# Machine Learning-Driven Discovery of Synergistic Protein Interactions Identifies ATL3 as a Putative Biomarker for Cancer Drug Response

**DOI:** 10.1101/2025.11.05.686871

**Authors:** Priya Ramarao-Milne, Anubhav Kaphle, Roc Reguant, Anne Klein, Michael Kuiper, Brendan Hosking, Julika Wenzel, Hawlader Al-Mamun, Lujain Elazab, Qing Zhong, Roger Reddel, Laurence Wilson, Yatish Jain, Natalie A. Twine, Denis C. Bauer

**Affiliations:** Australian e-Health Research Centre, Commonwealth Scientific and Industrial Research Organisation, New South Wales, Sydney, Australia; Department of Biomedical Sciences, Macquarie University, New South Wales, Sydney, Australia; Data 61, Commonwealth Scientific and Industrial Research Organisation, Canberra, ACT, Australia; Applied BioSciences, Faculty of Science and Engineering, Macquarie University, New South Wales, Sydney, Australia; ProCan®, Children’s Medical Research Institute, Faculty of Medicine and Health, University of Sydney, Westmead, New South Wales, Australia; University of Adelaide, Australian Institute for Machine Learning, Adelaide, Australia

**Author notes:** Authors contributed equally (Natalie A. Twine, Denis C. Bauer).

**Keywords:** machine learning, cancer, proteomics, synthetic lethality, drug susceptibility

## Abstract

**Background:** Resistance to targeted molecular therapies—both primary and acquired—remains a major obstacle to effective cancer treatment. Despite extensive research, the molecular determinants of treatment resistance are still incompletely understood, underscoring the need to identify robust resistance drivers to improve therapeutic outcomes. Recently, the ProCan and Wellcome Sanger Institute released the world’s largest pan-cancer proteomic dataset to date, comprising 949 cancer cell lines treated with 625 anti-cancer agents. Despite progress in identifying single protein markers, scalable methods for detecting synergistic protein interactions driving drug susceptibility remain limited. Machine learning models such as random forests may offer a robust framework for capturing non-linear protein interactions across diverse cancer types.

**Results:** Our study presents *synerOmics*, a scalable framework for identifying putative synergistic protein interactions by leveraging parent–child co-occurrences in random forest regression trees to reduce the interaction search space. We validated our approach using simulated data and two independent cancer proteomic datasets, identifying both pan-cancer and breast cancer-specific markers associated with drug susceptibility. Synergistic interactions that consistently replicated across datasets were enriched for endoplasmic reticulum stress pathways. Notably, shared targets included established sensitivity markers for tyrosine kinase inhibitors (TKIs) and revealed a resistance-associated network centred on ATL3, which demonstrated both prognostic and predictive relevance.

**Conclusions:** Applying synerOmics to the largest cancer cell line proteomic dataset to date, we identified ATL3 as a strong candidate biomarker for lapatinib resistance in breast cancer, with superior predictive performance compared to the current gold standard sensitivity biomarker, ERBB2.

**Highlights:**

- We present a machine learning framework for drug response biomarker discovery, leveraging the largest proteomic cancer cell line dataset, ProCan-DepMapSanger
- ATL3 is proposed as a candidate predictive biomarker for lapatinib resistance in breast cancer
- Elevated expression of a novel resistance-conferring subcluster centered on ATL3 is associated with poor prognosis across 11 TCGA cancer types
- Baseline drug resistance may be broadly driven by stress pathways, with ER stress and protein ATL3 implicated in ER-phagy in response to TKIs.

## Background

Primary and acquired resistance to targeted molecular therapies remain major obstacles to effective cancer treatment. Identifying reliable predictive biomarkers of drug susceptibility is critical to the advancement of precision oncology, however the molecular determinants of resistance are often challenging to identify, primarily because drug response mechanisms are multifactorial and influenced by the activity of multiple effectors. While genome-wide screens have revealed ∼2,000 pan-cancer essential genes^1^, emerging evidence suggests that drug susceptibility is shaped by complex, polygenic interactions more akin to complex traits and diseases^2^. In complex diseases, it has been observed that the effect of one or more genes can modulate the expression of another gene’s phenotype^3^, a phenomenon known as epistasis^1^. Such phenomena have also been detected in the context of cancer, involving “synergistic alterations” which tend to co-occur in a tumour, as well as “redundant” and “antagonistic” alterations which are mutually exclusive^4^. For example, in some cancers such as Lynch syndrome and hereditary breast and ovarian cancer syndrome, the genetic background of an individual can greatly influence the disease phenotype exerted by Tier 1 (high-confidence, clinically actionable) genetic variants^5^. This highlights the complexity of genetic influences in monogenic diseases where single, highly damaging mutations can affect disease risk, however multiple other weak effects may contribute to a “polygenic risk score” of sorts, leading to a continuum of disease phenotypes^5^.

Similarly, key molecular and cellular processes in cancer often arise from the combinatorial actions of multiple gene products, indicating that additive models may be insufficient to capture the complex, non-linear interactions underlying drug response^2^. Genome-wide screens studying single gene knockouts may deem certain genes as non-essential when screened in isolation and may be masking “synthetic essential” genes – whereby simultaneous perturbation of these genes diminishes cell viability and results in cell death, an effect called “synthetic lethality”. As such, a key strategy to understanding these synthetic lethal effects is to perform combinatorial knockout screens such as RNA interference (RNAi)^6^ or clustered regularly interspaced short palindromic repeats (CRISPR) screens^7^ followed by mathematical modelling to capture the interplay of molecular networks^8^. For example, the BRCA1–PARP1 synthetic lethality exemplifies such complexity: its penetrance is modulated by additional factors like TP53BP1^9^, RF1^10^ and PARG^5^, forming a network of co-dependencies --or “synergies”--that influence drug response. This highlights the need for a systems-level understanding of molecular effectors that exert synergistic, rather than additive, influences on drug sensitivity and resistance^9^.

Synergistic interactions have been shown to drive both drug sensitivity and resistance, presenting new opportunities for combination therapies. In fact, a recent study identified that targeting 2 regulators in acute lymphoblastic leukemia leads to significantly enhanced cell death compared with targeting either gene alone, demonstrating a synergistic effect^11^. Similarly, synergistic functions between paralogous genes have been shown to drive synthetic lethality in cancer, with co-inhibition of specific paralogs leading to markedly reduced cancer cell viability^12^. Landmark studies by the Sanger and Broad Institutes have resulted in the Cancer Dependency Map (DepMap)^13^, a large-scale resource cataloguing gene essentiality across diverse cancer types. While these studies have significantly advanced the identification of single-gene dependencies, they remain constrained by reliance on prior biological knowledge^14^ and do not account for combinatorial genetic interactions. As a result, they may overlook context-specific vulnerabilities that emerge from higher-order or synergistic effects in response to anticancer drugs.

A major challenge in studying synergistic dependencies lies in the immense combinatorial search space. For example, the number of *k*-way interactions tested for *n* number of genes 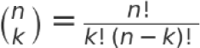 is given by and grows exponentially, scaling to over 6.6 quadrillion possible four-gene combinations. Therefore, *in-silico* approaches are required to systematically prioritise targets that may have putative synergistic relationships. *In-silico* approaches can provide heuristic alternatives to approximate how combined protein expression exerts an effect on drug susceptibility. Most previous work has explored the complexity of drug synergies^15^ in the context of identifying ideal combination therapies, with most methods involving mathematical models^16^, network analyses^17^ and ensemble methods^18^.

Some tools in this space include VIPER^20^, which specifically infers protein activity from gene expression data to detect synergies, and OPTICON^21^, which is based on the network controllability theory, however both require gene expression data as input. Other existing approaches for identifying interacting features utilise explainability algorithms —such as prototype networks^22^, local interpretable model-agnostic explanations (LIME)^23^, and Shapley additive explanations (SHAP)^24^— and primarily focus on explaining classification decisions. Importantly, these methods still do not adequately capture complex, multifactorial interactions between features which can be crucial to understanding molecular mechanisms in cancer networks. Additionally, these approaches rely on prior knowledge and existing databases^14^, limiting their capacity to uncover novel combinatorial vulnerabilities.

Machine learning and random forests, on the other hand, has shown promise for identifying novel and complex, non-linear relationships between features. Interestingly, a recent paper implemented deep exploration of random forests to identify interacting features between 16 variables relevant to the efficacy of lung nanoparticle targeting^25^. However, the deep exploration strategy implemented in this method becomes computationally infeasible when applied to high-dimensional omics datasets, which often contain thousands to millions of features. These limitations collectively underscore the need for a scalable, data-driven framework capable of uncovering novel synergistic interactions without relying on predefined networks or expression data.

To address this gap, we present synerOmics (**Figure 1**) which leverages random forests to capture pairwise synergistic interactions between continuous variables, such as proteomic, gene expression and other ‘omics data. Unlike previous approaches^25^, synerOmics can evaluate interactions at a genome-wide scale. Operating under the assumption that interacting features will occur in close proximity in random forest trees^25,26^, we reduce the search space of putative synergistic protein interactions by quantifying the occurrence of protein pairs as parent child nodes in the trees.

**Figure 1:**
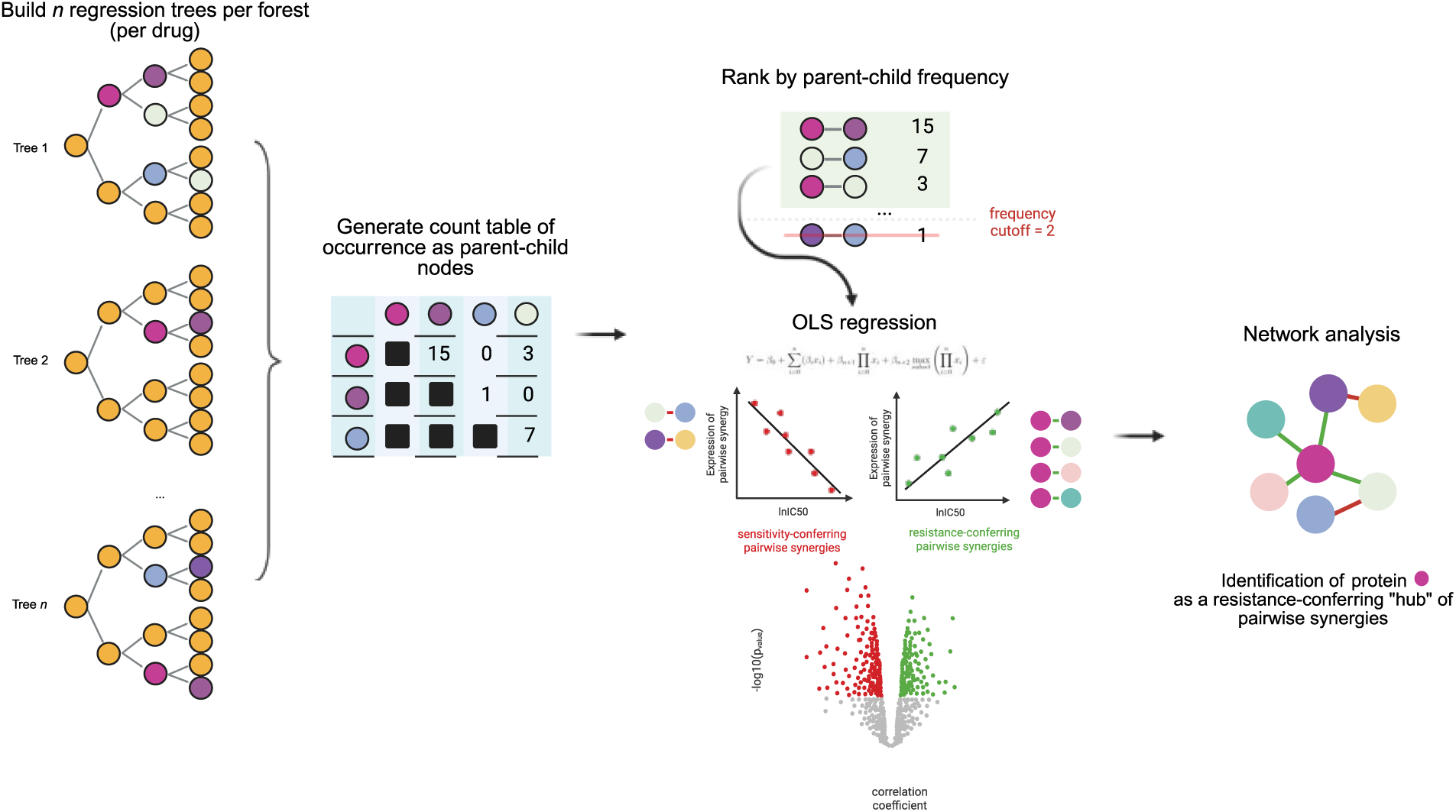
Graphical representation of research approach. synerOmics utilizes a 2-step approach: First, a random forest is trained on all features (proteins) in each dataset and a count matrix is then generated to determine how many times proteins appear together in parent-child orientation within the random forest. Any interaction that appears 2 or more times are retained for interaction testing. The second step to synerOmics involves running an OLS model to identify protein combinations which are synergistically and significantly correlated with drug sensitivity and resistance. After identifying common protein-pairs in both real-world datasets for each drug, we performed network analysis on these proteins to identify hub proteins. For the synthetic dataset we assessed the efficacy of synerOmics to detect known ‘truth’ interaction pairs and triplets in different interaction models.

In this paper we focus on proteomic data as increasing evidence shows that correlation between protein expression and drug susceptibility phenotypes is stronger than correlation with transcriptomic and genomic data^27–29^. We first demonstrate the efficacy our approach on simulated synthetic proteomic data, mimicking several interaction models adapted from previous studies^26,30^. We then apply our method to a pan-cancer cell line proteomic dataset^28^. These proteins may engage in direct interactions or participate in shared pathways^14^, suggesting that we can leverage pairwise synergistic relationships to build networks that may expose highly connected sensitivity or resistance hubs^31,32^. We therefore integrate synerOmics with network analysis to identify synergistic relationships between proteins that underlie drug susceptibility, validating these relationships in an independent breast cancer dataset^33^. Notably, our method operates independently of prior knowledge about features or drug targets, enabling the discovery of potentially novel relationships and alternative therapeutic targets.

## Results

### synerOmics captures synergies in synthetic proteomic data

To evaluate whether synerOmics effectively identifies synergistic pairs, we generated ten synthetic datasets featuring simulated drug IC_50_ phenotypes **(Supplementary Table S1, Supplementary Figures S0 - S2)**. We adapted 9 gene interaction models from Wright *et al.*^26^, with a primary focus on models containing synergistic interactions. Each dataset was designed to reflect either a *simple* phenotype - driven by a single ground truth pair from one interaction model, or a *complex* phenotype - driven by up to four ground truth pairs sampled from various models **(Supplementary Table S1)**. To simulate varying levels of biological noise commonly observed in real-world data, we introduced five heritability levels ranging from 10% to 100%, with 100% representing no noise.

We first tested *step 1* of synerOmics, which screens for potential interactions by fitting regression trees to the data and extracting all parent-child pairs (Figure 2A). We computed their occurrence counts across trees, retaining only parent-child pairs that occur more than twice (cutoff = 2). We then plotted the weighted detection rate of true interacting pairs (line graph in Figure 2B), which defines whether the truth set passed the cutoff threshold of 2. The weighted detection rate is calculated as a weighted average of detection rates across different levels of heritability. In this method, each detection rate is weighted by the inverse of its corresponding heritability value (i.e., weight = 1 / heritability). This weighting scheme gives greater importance to detection performance at lower heritability levels, where the signal is weaker and noise is higher. We then ranked the pairs according to frequency from highest to lowest and report the median count across 10 simulation runs, thus comparing the counts of our truth set against all extracted pairs.

**Figure 2:**
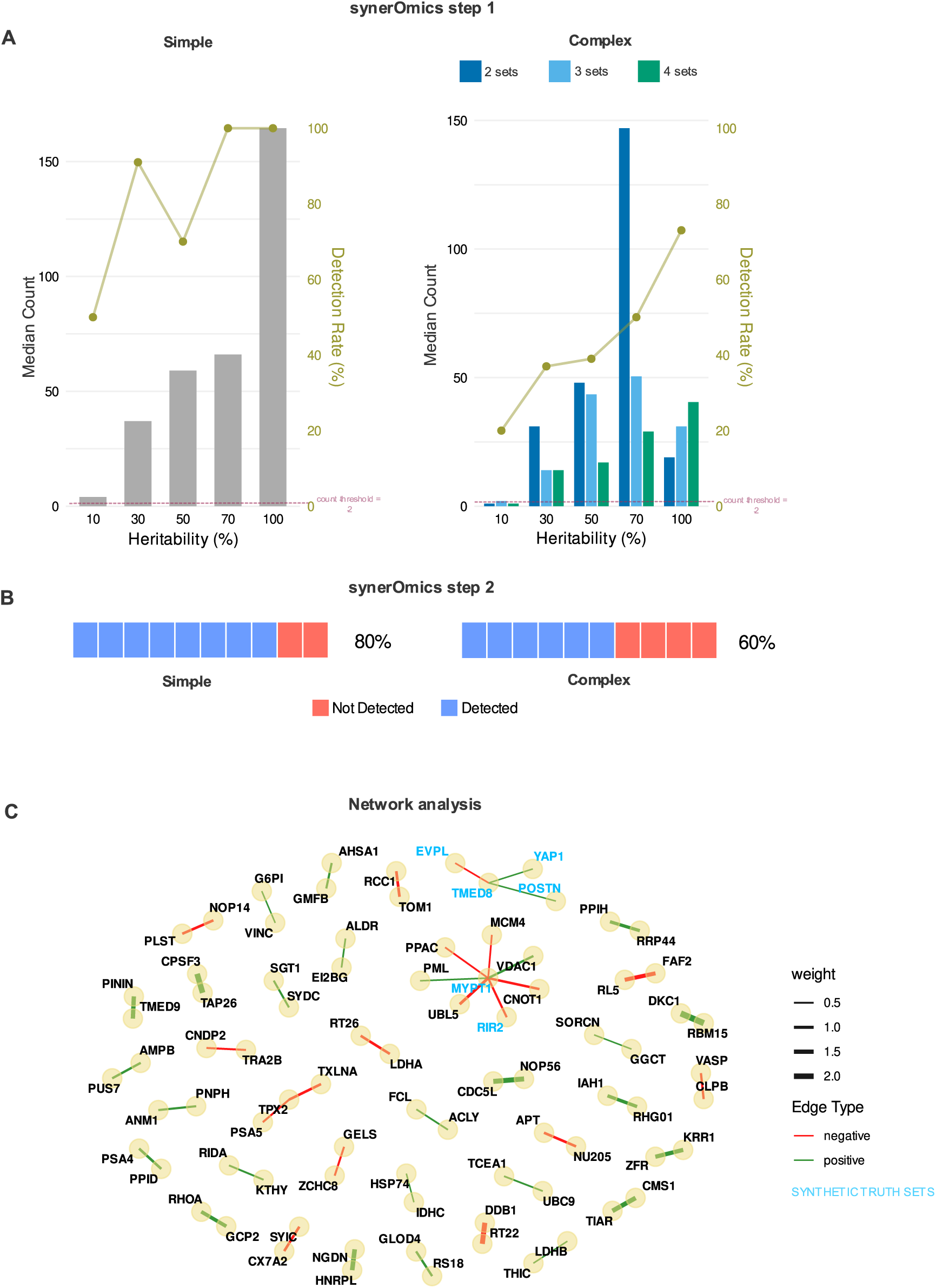
Performance of the random forest model in identifying true interacting protein pairs across synthetic datasets. **A**, Median counts of true interacting protein pairs are shown across different interaction models and heritability values (10%, 30%, 50%, 70%, 100%). For each model, we extracted parent-child node pairs from all 1500 trees generated by R’s Ranger package, counted their occurrences, and determined the dense rank of the true simulated interacting pairs based on these counts. Results are based on 10 synthetic datasets for each model-heritability combination, generated using the Goncalves et al. protein data matrix (900 x 6700). Higher counts indicate better detection capability of the random forest approach (mtry=500), with rank 1 representing optimal detection of the true protein interactions through their co-occurrence patterns. Line plot represents true variable pairs detection rate (%) trajectory for each interaction model across the heritability spectrum. The overall detection is calculated based on a weighted average of the detection rates across heritability values, emphasising larger weights for successful detection at lower heritability values. **B,** Detection rate of synerOmics step 2. A truth set is considered detected if it is identified as significant (adjusted p-value < 0.01) following regression analysis. **C,** Representative network plot of synthetic dataset analysis. Proteins coloured in blue text represent synthetic truth sets.

Figure 2A shows that for “simple” phenotypes, *step 1* of synerOmics detects 100% of the synergistic truth sets at a heritability of 70% and higher (low noise) and has an overall detection rate of 66.6% across the heritability spectrum. For “complex” phenotypes (interactions containing up to 4 truth sets of different interaction models, S1: Synthetic data experiments), the overall detection rate for synergistic pairs drops to 29.89% though our targeted interaction type (synergistic) has the third highest detection rate **(Supplementary Figure S3, Supplementary Figure S4)**. However, we observed that “Interaction only” and “Modifier protein” interaction models have lower detection rates especially with smaller effect sizes (3.40% and 9.65% respectively) **(Supplementary Figure S4)**. These findings indicate that synerOmics effectively identifies potential protein synergies, though its sensitivity may be reduced in scenarios with weak interaction effects, consistent with previous studies^26,34^.

Next, we assessed *step 2* of synerOmics: identify interactions that contribute beyond their individual (marginal) effects (Figure 1A). Here we employed ordinary least squares (OLS) regression to identify protein pairs for which the combined expression exerts a greater-than-additive effect on drug sensitivity or resistance. In our study, we define a resistance biomarker as having an expression level that is positively correlated with drug resistance, as indicated by a positive coefficient in an OLS regression model predicting log-transformed IC_50_ (lnIC_50_) values. Conversely, a negative coefficient suggests a sensitivity biomarker, reflecting increased drug sensitivity with higher expression. A protein pair that is identified during the first step as potentially synergistic is confirmed as such if the adjusted *p*-value (Benjamini–Hochberg procedure) for its corresponding interaction term in the OLS model is < 0.01. We generated an additional 10 datasets (simple and complex) at 70% heritability to quantify synerOmics’ capability to identify the synergistic truth sets. We observed an 80% detection rate for the simple phenotype synthetic datasets, and 60% for complex datasets (Figure 2B). Taken together, this confirms that if an interaction is successfully captured by *step 1* of synerOmics, *step 2* effectively distinguishes between marginal and synergistic effects, as *step 1* can also output marginally acting proteins as potential interaction partners in our approach due to the inherent properties of random forests.

Lastly, we assessed synerOmics’ ability to construct networks of synergistic proteins and identify “hubs” that have concentrated resistance or sensitivity pairwise synergies. For this we generate a dataset of complex phenotypes comprising of clusters of 5 to 10 synergistic pairs. To visualize the results, we use Cytoscape to construct a network by assigning the first protein in each synergistic pair as the source node and the second protein as the target edge. Figure 2C shows a representative example from one cluster, in which synerOmics successfully identified 4 out of 7 known synergistic pairs (57%). Here, network analysis identifies MYPT1 and TMED8 as hub proteins, exhibiting the highest betweenness, centrality and edge count (Figure 2C). This suggests that building a network of our putative pairwise synergies may reveal hub features that play a central role in mediating synergistic interactions.

Overall, synerOmics detects synergistic interactions across diverse synthetic interaction models, enabling the reconstruction of synergy networks, demonstrating its potential for gaining new insights from real world data.

### Shared pairwise synergies reflect drug class- and subclass-mechanism of action

We then applied synerOmics to the ProCan-DepMapSanger dataset, currently the most comprehensive proteomic dataset of human cancer cell lines, along with matched drug susceptibility data (949 cell lines and 8498 proteins). To validate our results, we used an independent dataset^33^, consisting of 76 breast cancer cell lines treated with 90 anti-cancer compounds, and proteomic data for 6091 proteins overlapping with the ProCan-DepMapSanger dataset. We selected a breast cancer-specific dataset for validation to account for tissue-specific biological differences that may influence drug susceptibility. Therefore, shared pairwise synergies that replicate in both datasets may reflect biologically relevant interactions within the context of breast cancer.

First, we interrogated all shared pairwise synergies common to both datasets. We only considered pairwise synergies if they were significantly associated with sensitivity or resistance in *both datasets* for the *same drug*, to determine if there were any synergies that were drug-class or drug-specific.

Overall, we identified 392 pairwise synergies which fell into 4 main drug classes: tyrosine kinase inhibitors (342 hits, 9 drugs), DNA inhibitors (16 hits, 5 drugs), microtubule inhibitors (11 hits, 3 drugs) and histone deacetylase (HDAC) inhibitors (11 hits, 1 drug) and other (12 hits, 11 drugs). **(Supplementary Tables S2 – S6)**. Of the 392 pairwise synergies across all drug class, 183 were unique hits. **Table 1** shows the most commonly occurring interactions overlapping between the datasets.

**Table 1:** This Table illustrates significant shared interactions, each occurring more than 5 times in both datasets. ProCan (S) and ProCan (R) columns denote number of occurrences in the ProCan-DepMapSanger dataset as sensitivity and resistance synergies respectively, and Validation (S) and Validation (R) columns denote number of occurrences in the validation dataset as sensitivity and resistance synergies. Synergies featuring a common drug in both datasets with a significant interaction are detailed under Common Drugs. The total number of occurrences across both datasets is summarised under the Total column. Interactions associated with at least one tyrosine kinase inhibitor are italicised.

| Pair | ProCan | ProCan | Validation | Validation | Total | Common Drugs | Drug class |
| --- | --- | --- | --- | --- | --- | --- | --- |
|  | (S) | (R) | (S) | (R) |  |  |  |
| *EGFR \* PAIRBP1* | 0 | 7 | 0 | 2 | 9 | *Afatinib* | *Tyrosine Kinase Inhibitors* |
| *ADAM9 \* PAIRBP1* | 0 | 7 | 0 | 2 | 9 | Paclitaxel, CHEMBL202721 | Microtubule Inhibitor |
| *TBL3 \* PAIRBP1* | 0 | 6 | 0 | 4 | 10 | Vinorelbine, Gemcitabine | DNA Inhibitor, Microtubule Inhibitor |
| *ESYT2 \* LMNA* | 0 | 6 | 0 | 2 | 8 | *Sorafenib, Vorinostat* | *Tyrosine Kinase Inhibitors* |
| *ESYT2 \* B2CL1* | 0 | 6 | 0 | 2 | 8 | *Sorafenib, Erlotinib* | *Tyrosine Kinase Inhibitors* |
| *MYO1B \* PAIRBP1* | 0 | 5 | 0 | 4 | 9 | Docetaxel | Microtubule Inhibitor |
| *MYO1C \* LMNA* | 0 | 5 | 0 | 3 | 8 | *Sorafenib, Oxaliplatin, Vorinostat* | *DNA Inhibitor, Tyrosine Kinase Inhibitor* |
| *PAIRBP1 \* WDR36* | 0 | 5 | 0 | 3 | 8 | Epirubicin | DNA Inhibitor |
| *PAIRBP1 \* MBB1A* | 0 | 5 | 0 | 2 | 7 | *Pevonedistat, Crizotinib* | *NAE Inhibitor, Tyrosine Kinase Inhibitor* |
| *EGFR \* VIME* | 0 | 4 | 0 | 2 | 6 | *Bosutinib, Gefitinib* | *Tyrosine Kinase Inhibitors* |
| *SLC3A2(4F2) \* LAMB3* | 5 | 0 | 4 | 0 | 9 | *Afatinib, Erlotinib, Gefitinib* | *Tyrosine Kinase Inhibitors* |
| *LAD1 \* LAT1* | 6 | 0 | 2 | 0 | 8 | *Erlotinib, Gefitinib* | *Tyrosine Kinase Inhibitors* |
| *LMNA \* BUD31* | 5 | 0 | 3 | 0 | 8 | Carboplatin, Oxaliplatin | DNA Inhibitors |
| *LAT1 \* DSG2* | 5 | 0 | 3 | 0 | 8 | *Afatinib, Erlotinib, Gefitinib* | *Tyrosine Kinase Inhibitors* |
| *EGFR \* KC1A* | 5 | 0 | 2 | 0 | 7 | *Erlotinib, Gefitinib* | *Tyrosine Kinase Inhibitors* |
| *EGFR \* DSG2* | 5 | 0 | 2 | 0 | 7 | *Erlotinib, Gefitinib* | *Tyrosine Kinase Inhibitors* |
| *SLC3A2(4F2) \* PLAK* | 5 | 0 | 2 | 0 | 7 | *Erlotinib, Gefitinib* | *Tyrosine Kinase Inhibitors* |
| *K1C15 \* LAT1* | 5 | 0 | 2 | 0 | 7 | *Erlotinib, Gefitinib* | *Tyrosine Kinase Inhibitors* |
| *LAD1 \* EGFR* | 4 | 0 | 3 | 0 | 7 | *Afatinib, Gefitinib* | *Tyrosine Kinase Inhibitors* |
| *EGFR \* PLAK* | 4 | 0 | 3 | 0 | 7 | *Gefitinib* | *Tyrosine Kinase Inhibitor* |
| *LAD1 \* SLC3A2(4F2)* | 4 | 0 | 2 | 0 | 6 | *Gefitinib* | *Tyrosine Kinase Inhibitor* |
| *EGFR \* SLC3A2(4F2)* | 4 | 0 | 2 | 0 | 6 | *Gefitinib* | *Tyrosine Kinase Inhibitor* |
| *EGFR \* K1C13* | 4 | 0 | 2 | 0 | 6 | *Erlotinib, Gefitinib* | *Tyrosine Kinase Inhibitors* |
| *EGFR \* K1C15* | 4 | 0 | 2 | 0 | 6 | *Erlotinib, Gefitinib* | *Tyrosine Kinase Inhibitors* |
| *EGFR \* S10A2* | 4 | 0 | 2 | 0 | 6 | *Erlotinib, Gefitinib* | *Tyrosine Kinase Inhibitors* |
| *EGFR \* LAT1* | 4 | 0 | 2 | 0 | 6 | *Erlotinib, Gefitinib* | *Tyrosine Kinase Inhibitors* |

Pathway analysis of these shared synergies showed that “regulation of apoptotic signalling pathway”, “protein localisation to cell periphery” and “response to oxidative stress” (Figure 3) as the most enriched pathways for our resistance synergies across both datasets. For synergies with the strongest effects (coefficient > 0.5 and <-0.5) we find associations with oxidative stress **(Supplementary Figure S6)**, specifically “ribosome biogenesis” and “cellular response to chemical stress”. Oxidative stress and disruptions in ribosome biogenesis have been shown to induce endoplasmic reticulum (ER) stress across multiple disease contexts^35^. ER stress is central to the development of drug resistance, as cells often rely on adaptive stress responses, including those activated by ER stress, to survive under adverse conditions such as those induced by chemotherapeutic agents^36,37^. This finding is consistent with reports within the literature that suggest that resistance to oxidative and ER induced stress-mediated cell death is linked to high proliferation rates and general drug resistance across most drug classes^38^.

**Figure 3:**
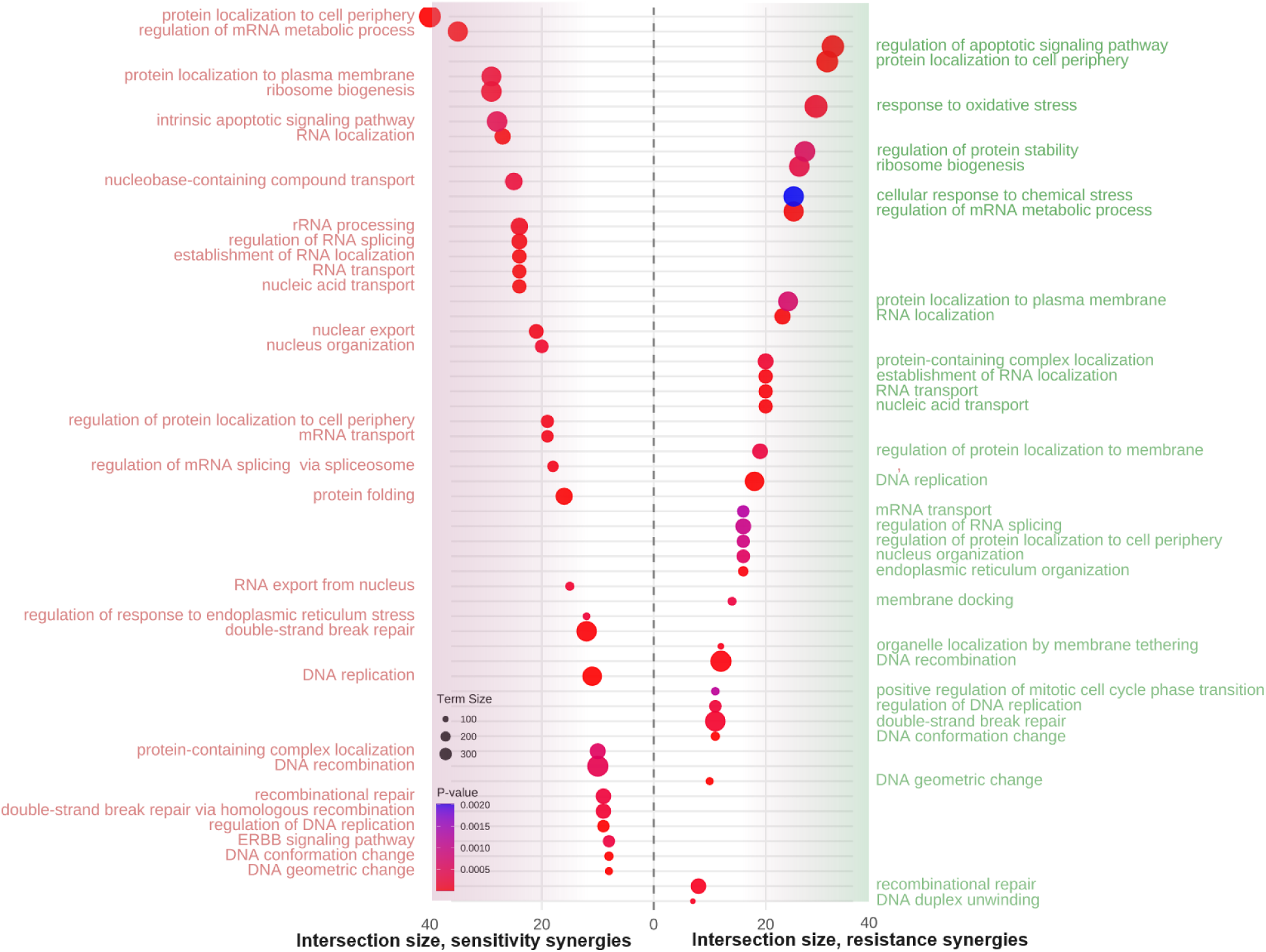
Pathway analysis of shared synergies. goProfiler analysis of shared synergies in both datasets, coloured by enrichment for resistance synergies (green text) and sensitivity synergies (red text). Pathways to the left of the dotted line represent pathway enrichment of sensitivity synergies, while pathways to the left of the dotted line represent pathway enrichment for resistance synergies. Circle sizes represent overall size of the pathway term, and colour represents p-value.

To gain a deeper understanding of the most commonly occurring pairwise synergies, we interrogated those at 5 or more occurrences (26 synergies). We observed that the majority of common hits belong to the tyrosine kinase inhibitor (TKI) drug class (21/26, represented by italicised text in **Table 1**).

Specifically, we identified EGFR (epidermal growth factor receptor) as a sensitivity biomarker for tyrosine kinase inhibitors such as afatinib, erlotinib and gefitinib, occurring 9/16 times (**Table 1, Supplementary Table S3)**. This is a well-established sensitivity marker, where activating EGFR mutations in a patient’s tumour are highly predictive of response to EGFR-targeting tyrosine kinase inhibitors such as erlotinib, gefitinib, and osimertinib^39^. EGFR occurs in two pairwise synergies (EGFR*DSG2 and EGFR*SLC3A2) for gefitinib and erlotinib, with both proteins already known to be part of the EGFR interactome^40^, thus validating known interactions.

We observed that drugs targeting the same pathway or sharing similar mechanisms of action exhibit overlapping synergy profiles **(Supplementary Figure S7A)**. MEK inhibitors trametinib and selumetinib shared the most extensive overlap with 38 common synergistic partners, while EGFR inhibitors erlotinib and gefitinib shared 20 synergies and dual EGFR/ERBB inhibitors afatinib and lapatinib shared 10 synergies. Notably, 7 of the 20 shared synergies between erlotinib and gefitinib involved EGFR, consistent with their common target specificity. Importantly, ERBB2 emerged as a sensitivity biomarker exclusively in the context of dual EGFR-ERBB inhibitors and was absent from all other drug classes **(Supplementary Figure S7B)**.

These observations indicate that drugs with shared mechanistic properties or targets may generate characteristic synergy landscapes that reflect their underlying target engagement patterns. Therefore, this mechanistic clustering of synergies may reveal novel latent therapeutic vulnerabilities which can be leveraged for experimental validation.

### Pairwise synergies in the tyrosine kinase inhibitor drug class converge on a novel resistance-conferring subcluster

We next sought to investigate deeper as to whether drugs with shared targets or mechanisms of action exhibit similar synergy profiles. TKIs were the most frequently represented drug class in our results, with 73% of all resistance-associated synergies and 94.4% of all sensitivity-associated synergies involving compounds from this class (Figure 4A). STRING’s Fisher’s exact test confirmed significant network enrichment (p < 1.0e-16; **Supplementary Table S11**), indicating more interactions than expected by chance. Pathway analysis further confirmed the drug mechanism of action with ‘cell adhesion molecule binding’, ‘cell junction reorganisation’, and ‘signalling by receptor tyrosine kinases’ **(Supplementary Table S12)**.

**Figure 4.**
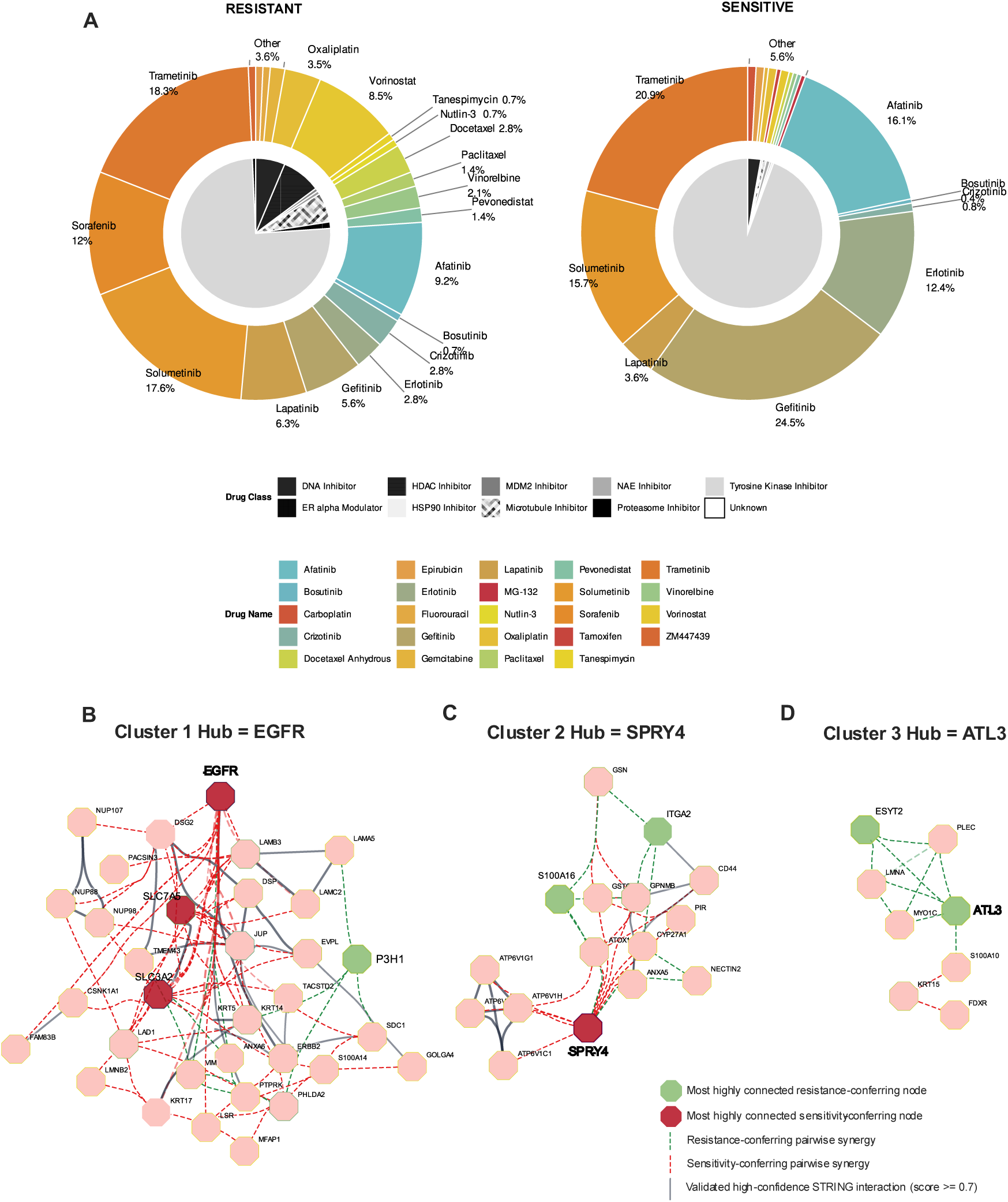
Pie chart representing drug class and drug name of shared hits. Inner chart in grayscale represents drug class, coloured doughnuts represent drugs. MCL clustering of significant pairwise interactions for Tyrosine Kinase Inhibitor networks centred around nodes with highest edge count and betweenness centrality for three main clusters; **B,** Cluster 1: EGFR, **C,** Cluster 2: SPRY4 and **D,** Cluster 3: ATL3. Green dotted lines represent a positive correlation with IC50, representing a “resistance”-related synergy, whereas red dotted lines represent a negative correlation with IC50, representing a “sensitivity”-related synergy. Solid lines represent high-confidence validated STRING database interaction score (>0.7). For clarity, only nodes with degree >= 3 are shown. Filled red nodes are top candidates for highest connected sensitivity nodes, whereas filled green nodes represent candidate resistance nodes with highest degree and betweenness centrality. Hubs are represented by bold black font.

To further characterise TKI-associated synergies, we analysed the network using Cytoscape **(Supplementary Tables S7–S10)** and applied the Markov Cluster Algorithm (MCL) to identify sub-networks within the clusters. Node importance was quantified using ‘betweenness centrality’ to assess communication flow, and ‘edge count’ to evaluate connectivity and potential hub roles in biological systems. MCL clustering revealed three major clusters (Figure 4B **– 4D)**.

Unsurprisingly, the largest cluster was centred around EGFR, the gold-standard clinical biomarker for TKI efficacy. EGFR had the highest betweenness centrality and edge count (Figure 4B) (all network analysis metrics are reported in **Supplementary Tables S7 – S10**). A potential alternate marker for this network may be SLC7A5 (Solute Carrier Family 7 Member 5), which is part of cluster with only sensitivity-conferring interactions. The second most connected network (Figure 4C) is centred around SPRY4, another well-known negative feedback regulator of the MAPK signalling pathway which is downstream of EGFR receptor activation. This cluster has most connections to other proteins representing sensitivity-related synergies. In fact, SPRY4 has been widely studied as an alternative target in the case of third generation TKI resistance^41,42^.

The third most highly connected network (Figure 4D) is a potentially novel TKI resistance cluster centering around ESYT2 and ATL3. ESYT2 knockout was recently shown to induce ER-stress mediated cell death in mice^43^. ATL3 was found to function as a receptor for a selective form of autophagy called ER-phagy, promoting tubular ER degradation during cell starvation^44^. ER stress induces the formation of ER tubular bodies (ER-TBs), and a recent paper has consolidated the important role of ATL3 in the formation of ER-TBs in SARS-CoV-2^45^. Cancer cells exhibiting elevated basal ER stress are broadly resistant to anticancer therapies^46^; however, pharmacological inhibition of ER stress can restore drug sensitivity^47^. Metastatic cancer cells that are inherently more resistant to high levels of ER stress have a selective advantage due to increased survival capabilities^47^. Other proteins in this novel cluster (PLEC, MYO1C and LMNA) have also been implicated in cancer to some degree, but there are no previous links to ER stress that have been consolidated. Additionally, none of the proteins in our novel resistance cluster have been previously associated with TKI resistance.

Taken together, ER stress pathway is re-iterated as important for breast cancer with TKI emerging as central drug-target. Specifically, we identify a novel resistance sub-cluster consisting of ATL3, ESYT2, LMNA, MYO1C and PLEC for predicting resistance to tyrosine kinase inhibitors.

### TK resistance synergies define R-stress module with prognostic and predictive potential

We then investigated the clinical significance of the novel resistance-conferring cluster around ATL3. We performed Kaplan-Meier survival analysis using the mean expression of the five genes in the resistance cluster (*ATL3*, *ESYT2*, *LMNA*, *MYO1C* and *PLEC*) **(Supplementary Figure S8, Supplementary Table S13)**. We used mRNA levels as a proxy for protein expression as there is currently no publicly available pan-cancer proteomic patient tumour data associated with survival. Progression-free survival was assessed using hazard ratios, with Kaplan–Meier curves generated via KM-Plotter^48^ to visualise survival outcomes.

Kaplan–Meier survival analysis revealed that the ATL3 resistance-conferring cluster was significantly associated with progression-free survival in 11 out of 18 cancer datasets (Figure 5A). The most significant associations were observed in liver hepatocellular carcinoma (p = 0.018), pancreatic ductal adenocarcinoma (PDAC) (p = 0.023), and ovarian cancer (p = 0.024), as shown in Figure 5B. To evaluate the robustness of these associations and account for potential background noise, we generated two control groups: (i) a “Random” set comprising five proteins selected at random, and (ii) four “Marginal-only” sets composed of five proteins individually associated with resistance but not involved in any significant pairwise synergies. As expected, the Random set yielded no significant hazard ratios, while the Marginal-only sets showed limited associations, with a maximum of seven cancer types reaching significance. This reduced performance is consistent with the independent effects of marginal genes, underscoring the added predictive value of synergistic interactions.

**Figure 5:**
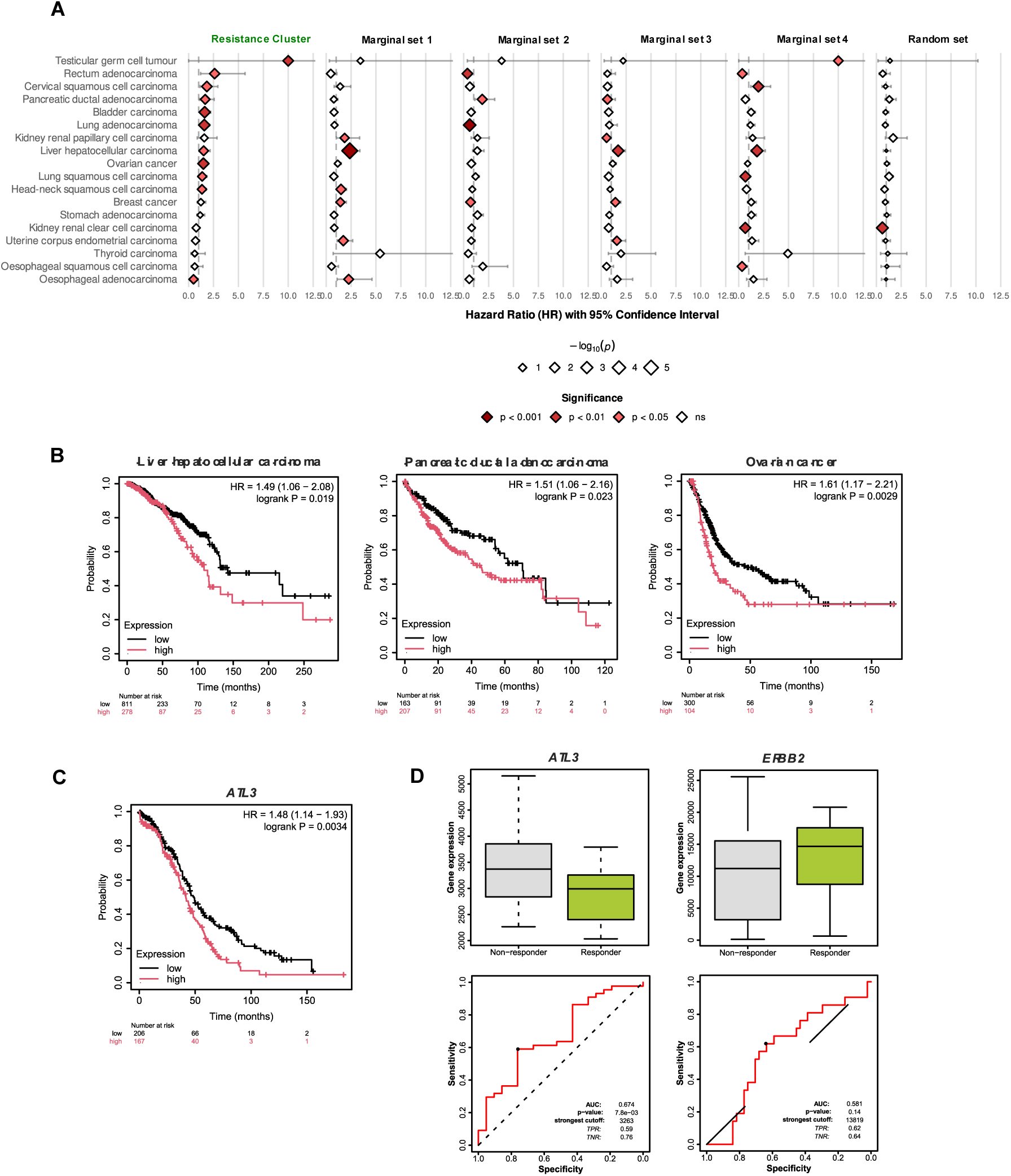
Clinical correlates of candidate biomarkers. **A**, Forest plots representing hazard ratios for our resistance cluster consisting of 5 genes, and four additional sets of five genes randomly sampled from corresponding proteins in our analysis that had marginal effects (positive coefficients) only, but no interactions and random subset of 5 genes. **B,** Representative Kaplan-Meier survival analysis of three TCGA mRNA datasets (liver hepatocellular carcinoma, pancreatic ductal adenocarcinoma and ovarian cancer) using gene signature (mean expression) of *ATL3*, *ESYT2*, *MYO1C*, *LMNA* and *PLEC*. Red asterisk * represents a significant association between worse survival and high expression of gene signature, black asterisk * represents a significant association between better survival and high expression of gene signature. **C,** Survival analysis of breast cancer TCGA protein dataset stratifying by *ATL3* only gene expression levels. **D,** Predictive biomarker analysis from ROCplot depicting predictive utility of ATL3 levels and ERBB2 levels in stratifying response to anti-HER2 therapy lapatinib. p-value is calculated from Mann-Whitney t-tests comparing mRNA levels of responders vs. non-responders. FC = fold-change

Single-gene biomarkers are preferred in clinical settings due to their diagnostic tractability, ease of validation, and compatibility with routine molecular profiling workflows. We therefore tested the ability of *ATL3* alone in predicting survival. Kaplan–Meier analysis demonstrated a significant association between *ATL3* expression and progression-free survival in breast cancer (p = 0.034, Figure 5C), despite the broader synergy cluster not reaching statistical significance in this context (p = 0.065, Figure 5A). To further evaluate the potential of ATL3 as a predictive biomarker for lapatinib response in breast cancer, we employed ROCplot^49^ to compare expression levels between responders and non-responders. Lapatinib, a dual EGFR and ERBB2 tyrosine kinase inhibitor, is approved for patients with HER2 (ERBB2) amplification. Notably, a significant difference in *ATL3* expression was observed (Mann–Whitney test, p = 0.0024), with responders exhibiting approximately 1.2-fold lower levels compared to non-responders (Figure 5D), supporting its potential utility as a candidate predictive biomarker for lapatinib resistance.

For comparison, we assessed ERBB2, the gene encoding the HER2 protein, which serves as a clinical inclusion criterion for lapatinib treatment in patients with HER2-amplified breast cancer. Although ERBB2 expression trended higher in responders (1.2-fold increase), this difference did not reach statistical significance (p = 0.14; AUC = 0.581). In contrast, ATL3 demonstrated superior discriminatory performance (AUC = 0.674, p = 0.0078), with improved specificity (true-negative rate = 0.76) at comparable sensitivity (true-positive rate = 0.59), suggesting enhanced utility in identifying non-responders.

Importantly, ATL3’s predictive performance improved when analysis was restricted to HER2-positive samples, yielding an AUC of 0.744 (p = 0.00044) and a true-negative rate of 0.82, whereas ERBB2 remained non-significant in this subgroup **(Supplementary Figure S10)**. These findings suggest that ATL3 may offer enhanced specificity within the clinically relevant HER2-positive population, reinforcing its potential as a negative predictive biomarker for lapatinib response. However, we acknowledge that the HER2-stratified analysis was limited by small sample size, and further validation in larger cohorts is warranted.

Taken together, these results support further investigation of *ATL3* as a putative biomarker for predicting non-response to anti-HER2 therapies such as lapatinib.

### Structural modelling highlights AT 3 as a potential candidate for therapeutic targeting

Lastly, we sought to investigate whether ATL3 possesses druggable structural characteristics that could influence lapatinib activity, and whether ATL3 may be a druggable candidate for resistance biomarker modulation. To this end, we applied AlphaFold3 and Fpocket to model and analyse protein– ligand interactions involving EGFR and ERBB2, lapatinib’s known targets, as well as the putative resistance biomarker ATL3.

We modelled the three possible ligand-protein pair combinations, lapatinib-EGFR (Figure 6A), lapatinib-ERBB2 (Figure 6B) and lapatinib-ATL3 (Figure 6C) separately. As expected, both the lapatinib–EGFR and lapatinib–ERBB2 complexes exhibited high model ranking scores (0.96 and 0.99, respectively) and low positional uncertainty (chain_pair_pae_min ≤ 1.22 and ≤ 1.19, respectively) (**Table 2**), consistent with stable and specific interactions. Comparatively, the lapatinib-ATL3 complex showed a moderate binding confidence (ranking score = 0.71; iPTM = 0.67) (**Table 2**) and elevated spatial uncertainty (chain_pair_pae_min up to 7.34) (**Table 2**), suggesting a weaker but still plausible interaction.

**Figure 6:**
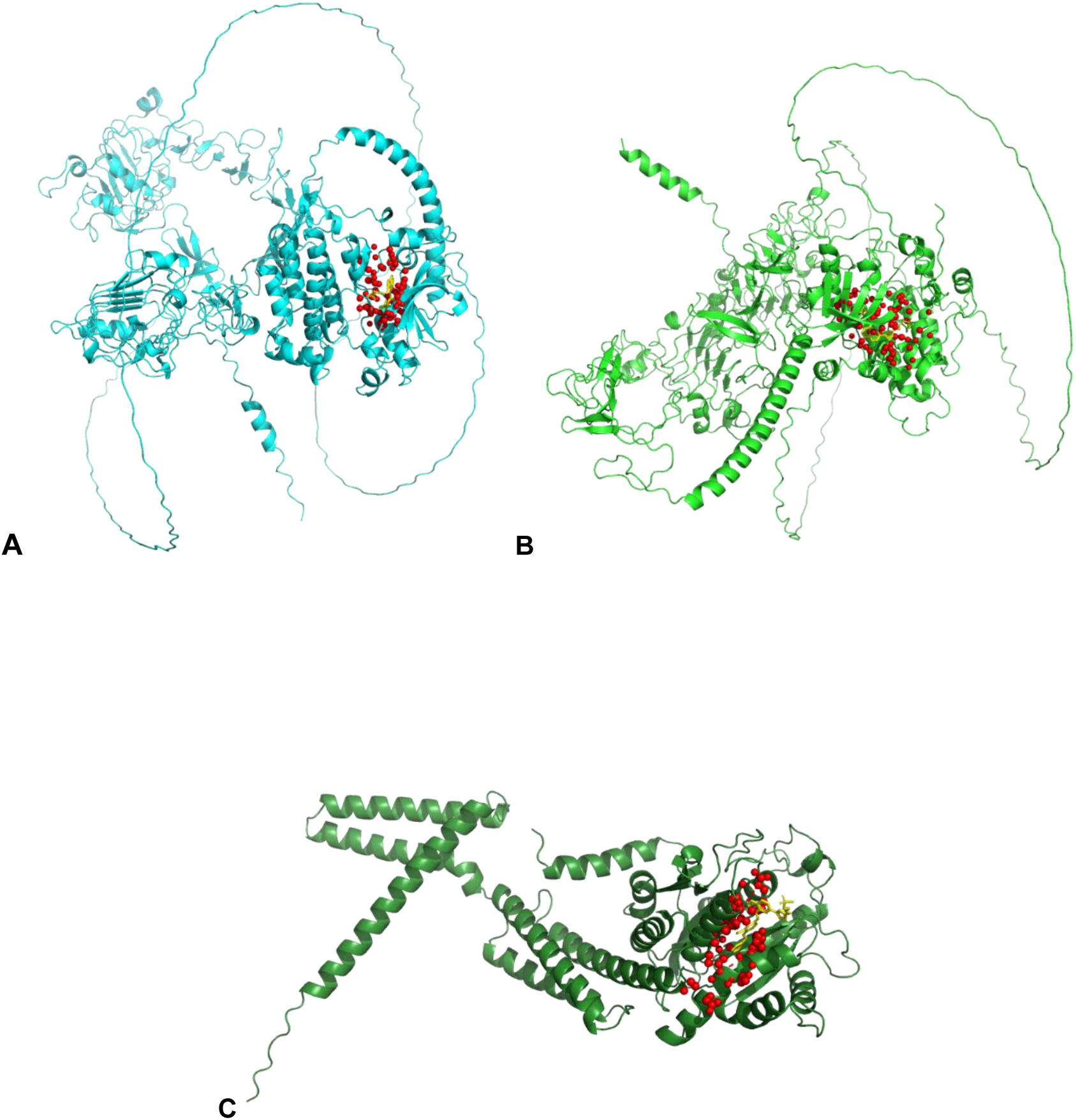
AlphaFold3 structures of **A,** lapatinib-EGFR complex, **B,** lapatinib-ERBB2 complex and **C,** lapatinib-ATL3 complex. The highest-confidence binding pockets (Pocket 1) are visualized as red spheres in all three structural models. The bound ligand is depicted as a yellow stick representation, highlighting its spatial orientation within the predicted pocket.

**Table 2:** details AlphaFold3 and AutoDock Vina metrics for lapatinib-EGFR, lapatinib-ERRB2 and lapatinib-ATL3.

| <u>AlphaFold3 Metric</u> | lapatinib-EGFR | lapatinib-ERBB2 | lapatinib-ATL3 |
| --- | --- | --- | --- |
| *iptm* | 0.96 | 0.96 | 0.67 |
| *ptm* | 0.44 | 0.56 | 0.7 |
| *ranking_score* | 0.96 | 0.99 | 0.71 |
| *chain_iptm* | [0.96, 0.96] | [0.96, 0.96] | [0.67, 0.67] |
| *chain_ptm* | [0.45, 0.65] | [0.56, 0.65] | [0.7, 0.51] |
| *chain_pair_iptm* | [[0.45, 0.96], [0.96, 0.65]] | [[0.56, 0.96], [0.96, 0.65]] | [[0.7, 0.67], [0.67, 0.51]] |
| *chain_pair_pae_min* | [[0.76, 1.09], [1.22, 0.77]] | [[0.76, 1.09], [1.19, 0.76]] | [[0.76, 5.91], [7.34, 0.77]] |
| *fraction_disordered* | 0.21 | 0.22 | 0.06 |
| *has_clash* | 0 | 0 | 0 |

| <u>AutoDock Vina Binding Score</u> | lapatinib-EGFR | lapatinib-ERBB2 | lapatinib-ATL3 |
| --- | --- | --- | --- |
| *energy optimised binding affinity*<br>(kcal/mol) | -10.143 | -10.47 | 18.799 |
| *raw pose binding affinity*<br>(kcal/mol) | -9.867 | -9.996 | 30.742 |

Given the moderate confidence and elevated spatial uncertainty observed in the lapatinib-ATL3 complex prediction, molecular docking analysis was performed to quantitatively assess binding feasibility across resistance-associated proteins. ERBB2 demonstrated the strongest binding affinity (−10.47 kcal/mol; energy-optimized vs. −9.996 kcal/mol raw pose) (**Table 2**), validating lapatinib’s established therapeutic mechanism. In stark contrast, ATL3 exhibited thermodynamically prohibitive binding energetics (+18.799 kcal/mol optimized; +30.742 kcal/mol raw pose) (**Table 2**), effectively ruling out direct competitive inhibition mechanisms. These energetically unfavourable interactions indicate that ATL3 does not directly interfere with lapatinib’s mechanism of action.

To evaluate broader therapeutic targeting potential of ATL3, we utilized Fpocket to systematically identify druggable binding pockets in each AlphaFold3-predicted model. We extracted residues in close proximity to the ligand lapatinib (within 4 Å) and computed two key metrics: the Fpocket score, which reflects the geometric and physicochemical suitability of a pocket for general ligand binding, and the Drug score, which specifically estimates the likelihood of a pocket accommodating drug-like molecules. These scores help rank pockets within each protein based on their structural stability and druggability. For EGFR (Figure 6A) and ERBB2 (Figure 6B), we observed high Fpocket scores (0.883, 0.617) and Drug scores (0.990, 0.963) respectively, consistent with well-defined, drug-compatible binding sites **(Supplementary Data D3 – D5)**. Interestingly, ATL3 (Figure 6C) exhibited an Fpocket score in between that of ERBB2 and EGFR (0.723), suggesting a less optimal pocket geometry, yet a similarly high Drug score (0.994), indicating that ATL3 harbors a structurally stable, druggable pocket. This is supported by the fact that the selected ATL3 pocket maximized overlap with highly conserved binding residues, identified as residues within 4 Å of lapatinib across ≥80% of AlphaFold3 models **(Supplementary Data D5)**.

Taken together, our analyses suggest that ATL3 contributes to TKI resistance via downstream signaling modulation rather than direct lapatinib sequestration, supporting its potential for combination therapy to circumvent resistance. Moreover, structural modelling and druggability analysis identify protein ATL3 as a therapeutically actionable target. These computational findings warrant experimental validation to clarify ATL3’s mechanistic role.

## Discussion

In this study, we present a novel approach to identify protein combinations that have a synergistic effect on drug susceptibility in cancer cell lines. We leverage a combination of ML and network analysis to identify novel protein pairs correlating with drug response, validating our method in multiple synthetic datasets and two independent real-world datasets. Our approach fills a gap to identify synergistic pairwise interactions in continuous input data types and continuous phenotypes, thus having application to a wide range of ‘omics datasets. Additionally, our systematic end-to-end approach, from machine learning algorithms to structural prediction and molecular docking, represents a hypothesis-free methodology for target prioritization, enabling identification of resistance mechanisms independent of existing biological assumptions.

Using synerOmics, we observe that cellular stress-response pathways, particularly those associated with endoplasmic reticulum (ER) stress, may play a role in mediating resistance to cancer therapeutics in breast cancer. HER2-positive (HER2+) breast cancers, accounting for approximately 15–20% of all breast cancer cases, are typically associated with aggressive tumour growth and poor patient prognosis^50^. Current HER2-targeted therapies exploit overexpression of ERBB2, the proto-oncogene encoding the HER2 receptor, to inhibit tumour invasiveness and metastasis. Insights from HER2+ gastric cancer suggest that endoplasmic reticulum (ER) stress may play a critical role in the development of trastuzumab resistance, highlighting a potentially conserved mechanism across HER2-expressing malignancies^51^, however in breast cancer, the biological mechanisms of resistance are unresolved^52^. Importantly, there are no clinically approved biomarkers currently exist to predict resistance to HER2-targeted therapies, limiting the ability to guide treatment decisions and manage therapeutic failure. In our work, ER stress was implicated as a resistance mechanism in breast cancer across multiple drug classes, with distinct molecular signatures predictive of therapeutic response of a broad range of anticancer agents. Notably, ATL3 emerged as a recurrent resistance-associated factor particularly within the TKI drug class, suggesting that its basal expression levels or associated interactome may influence drug susceptibility in cancer cells.

While ATL3 has not yet been directly implicated in tumorigenesis, its dysfunction is well-documented in neurodegenerative conditions, where loss-of-function mutations lead to aberrant ER morphology^53^. Specifically, pathogenic variants in ATL3 are causative of autosomal-dominant hereditary sensory neuropathy type 1F (HSN1F) through disruptions in the tubular ER network^54^, underscoring its physiological importance in ER homeostasis. ER stress is known to induce the formation of ER tubular bodies (ER-TBs), specialised membrane structures involved in stress adaptation^45^. Recent studies have identified ATL3 as a key structural determinant in ER-TB biogenesis, particularly in the context of SARS-CoV-2 infection^45^. In oncogenic settings, ATL3 has been linked to selective autophagy, functioning within a regulatory network of autophagy-related proteins^55^. In tumour contexts, ER-phagy can either support cancer cell survival by alleviating proteotoxic stress or promote cell death when excessively activated^56^, reflecting a dualistic role akin to that of the unfolded protein response (UPR) - where moderate ER stress fosters adaptation and growth, while unresolved stress leads to apoptosis^57^. Given that ATL3 has been implicated in ER morphology, ER-TB formation, and selective autophagy, it may function as a regulatory node within ER-phagy pathways in cancer.

Currently, only a limited number of clinically approved agents modulate ER stress-related pathways, such as carfilzomib^58^ and bortezomib^59^ in multiple myeloma – both proteasome inhibitors that activate the unfolded protein response either directly or indirectly. Importantly, their clinical use is limited by acquisition of resistance and substantial side effects in patients. Importantly, there are currently no clinically approved drugs targeting ER-phagy. Selective targeting of ATL3, a direct ER-phagy receptor may enable more precise modulation of ER stress signalling, potentially reducing off-target effects and improving therapeutic specificity. In clinical cohorts, analysis of the TCGA breast cancer dataset demonstrated that ATL3 expression was significantly higher in patients who did not respond to lapatinib treatment, particularly in the HER2+ subgroup, supporting its potential role as a predictive biomarker of therapeutic resistance, warranting further investigation *in-vitro*. Moreover, our druggability assessment indicates ATL3 possesses favourable binding pockets, supporting its candidacy for therapeutic development and *in vitro* validation. We note that while our binding affinity calculation did not support thermodynamically stable lapatinib-ATL3 complex, we cannot discount the possibility that any post-translational modifications and/or co-factors such as ions in real physiological context might stabilise the binding. Overall, our study suggests that ATL3 may be potential candidate for further investigation as a modulator of ER-phagy or stress pathways in breast cancer, with potential implications for overcoming therapy resistance and refining treatment strategies. While our *in silico* findings offer valuable mechanistic insights and support hypothesis generation, experimental validation is essential to clarify ATL3’s functional implications in the context of therapeutic resistance.

In addition to our work identifying novel cancer drug response biomarkers, we present a novel methodological framework demonstrating that synergistic protein pairs are preferentially represented as parent-child nodes in random forest models. Using simulated datasets encompassing diverse interaction types, we validate the robustness of our approach, while acknowledging that the models, originally designed for genetic epistasis, may not fully capture the complexity of protein interactions. Network construction based on pairwise synergies reveals key hub proteins associated with drug sensitivity and resistance and proteins potentially contributing to these phenotypes. This aligns with prior findings, such as the SynPathy study^60^, which demonstrated that drug combinations that have a synergistic effect on cell death often involve pathways that are in close proximity to one another in a protein interaction network. Our synerOmics framework effectively captures non-linear relationships, narrowing the search space for putative interactions even in the absence of marginal effects, and successfully identifies true interacting pairs. While interactions with weak marginal signals are ranked lower, deeper tree-parsing strategies may enhance interpretability. Although our current use of OLS regression is optimized for linear synergies, the framework is extensible to non-linear models (e.g., cubic, exponential, sigmoidal), which could further improve detection. Notably, these limitations do not affect our real-world cancer dataset analysis, where the primary objective was to identify synergistic interactions.

Taken together, we present synerOmics, an *in silico* framework for prioritising candidate biomarkers for *in vitro* validation, with potential for expansion through multi-omics integration to uncover cross-omic signatures of drug susceptibility. Notably, our findings identify ATL3 as a candidate resistance biomarker in lapatinib-treated breast cancer, addressing the current clinical gap by demonstrating superior predictive performance for non-responders, while ERBB2 remains the gold standard for identifying responders.

## Methods

### synerOmics method overview

Here, we adapt and extend the ML method developed by Al-Mamun *et al.,* 2022^61^, which utilises a random forest model, to identify pairwise and higher-order protein interactions that underlie drug response in our study (Figure 1). In the context of epistasis, Al-Mamun *et al* observed that interacting Single Nucleotide Polymorphism (SNP) markers appear together more frequently in a decision tree than non-interacting markers (e.g., as parent-child nodes). We have adapted this approach for continuous variables, thus extending its applicability to other data types, including proteomic and transcriptomic data. For readability, we herein describe the method as applied to proteomic data, as we utilized proteomic (real and synthetic) data in this study. SynerOmics identifies protein-protein relationships through a 2-step process: *(1)* random forest modelling to screen for potentially interacting variables, thereby reduce search space, followed by *(2)* linear regression with interaction terms to delineate protein pairs and their effect sizes impacting drug susceptibility. The key hypothesis assumed in the first step is that the proteins affecting phenotype through interaction would appear together more frequently (for example, as parent-child nodes) across decision trees than non-interacting proteins.

### synerOmics step 1: Quantification of parent-child appearances of features in regression trees

The first step of synerOmics involves reducing the search space of all possible pairwise interactions, via quantification of appearances of features as parent-child nodes. This step is conducted to capture features that occur with the highest frequency as parent-child nodes, under the premise that this approach will enrich our testing pool for putative synergistic interactions. Specifically, we applied synerOmics to our first real-world pan-cancer proteomic cell line dataset (ProCan-Sanger^28^). The dataset consists of a protein count table derived from data-independent acquisition mass spectrometry (DIA-MS) profiling baseline protein expression in 949 cancer cell lines, and drug sensitivity data (measured by the half-maximal inhibitory concentration (IC_50_)) for 625 anti-cancer drugs commonly used in the clinic. First, we trained a random forest model for each of the 625 cancer drugs, building regression trees for 6091 protein markers across the 949 cancer cell lines. We utilised the following parameters to build our regression trees; n_trees = 1500, min_node_size = 5, max_depth = 82, mtry = 500. Using information from the resultant regression trees, we generated a parent-child node frequency count matrix, to quantify how often different proteins appear as pairs, triplets and quadruplets in the regression trees. We utilised a cutoff of occurrences = 2 which means that the combination has to occur at least twice to be considered for the next step of synerOmics. Effectively, this approach selects for the pairs with the greatest potential to show synergistic interactions (Figure 2).

### synerOmics step 2: Ordinary Least Squares regression model

Next, we aimed to determine whether the selected pairwise synergies have a greater than additive effect on impacting drug susceptibility than each protein on its own. Specifically, we sought to determine whether the interaction effect was greater than the additive marginal effects for each protein. For this purpose, we employed an ordinary-least squares (OLS) model to evaluate the significance of identified synergistic interactions. We utilised the python package *statsmodels* OLS formula, whereby the beta coefficients correspond to the influence of a particular variable on the phenotype Y. In order to model the additional non-additive or “synergistic” effect of combined protein expression, our approach utilised a multiplicative epistasis model, wherein pairwise synergistic effects are modelled by β_3_. Interactions were characterised by:

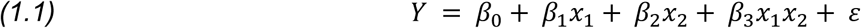

Where x_1_ and x_2_ represent the variables (proteins), β_1_ is the marginal effect for the first protein, β_2_ is the marginal effect for the second protein and β_3_ is the interaction effect. Y represents the phenotype, and E denotes the residual. Furthermore, to model higher order interactions, we extended our model to encompass:

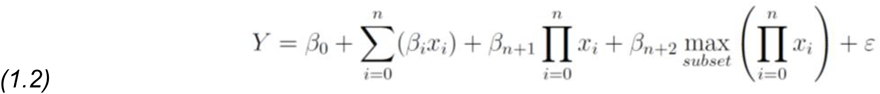

Which takes into account the maximum effect from the n-1th order. For example, in the context of a triplet synergy, we include the pairwise synergy with the highest effect in the regression model. This is to ensure that any observed non-additive effects are not merely attributable to a subset of the triplet synergy, and we have taken the maximum subset synergy to reduce the number of tests we need to conduct. In our analysis, a significant p-value for the interaction term allows us to reject the null hypothesis, which is that the interaction coefficient has no effect on effect on the phenotype, over the marginal effects. To account for multiple hypothesis test correction, the Benjamini-Hochberg method was applied to account for the false discovery rate (p-adjusted < 0.01 was deemed as a significant synergy). Only synergies that have a greater than additive effect on drug susceptibility were retained. Coefficients ≥ 0 denote a positive correlation with IC_50_ and are therefore considered drug resistance-conferring pairwise synergies, and coefficients ≤ 0 denote a negative correlation with IC_50_ and are considered drug sensitivity-conferring pairwise synergies.

### Validation of methodology with an independent dataset

In order to validate the interactions we detected in the original ProCan-Sanger dataset consisting of 949 cancer cell lines, we applied our synerOmics method to an independent dataset Sun *et al.,* 2023^33^. This dataset consisted of 76 breast cancer cell lines treated with 90 anti-cancer compounds, and proteomic data for 6091 proteins. All methodology was conducted as above for the primary dataset.

### Application of synerOmics to synthetic data

To test our hypothesis that features with a greater than additive effect occur more frequently as parent-child nodes than non-interacting features, we generated 10 synthetic datasets for each of 9 interaction models (detailed below) simulating drug IC_50_ phenotypes. We use the Goncalves et al.^28^ protein data matrix (cell-types = 900 x proteins=6700) for simulation to preserve any unknown confounding effects in the original data, to enhance the realism of our simulation.

Specifically, we simulated order-2 and order-3 protein interactions where applicable by randomly selecting proteins as *true variables* that contribute to the simulated phenotype. Interactions higher than order-3 were not consider in our testing regimen. We excluded proteins as *true variables* if they had any correlated partners in the data matrix based on the estimated Pearson’s correlation coefficient threshold of 0.70. This helps to avoid spurious associations due to correlation structures in the data. Where applicable, marginal effect sizes were fixed at −1 or 1, uniformly chosen each time.

Interaction effect sizes were sampled from a standard Gaussian distribution (with standard deviation = absolute value of (marginal effect size/3)). Gaussian noise was added to the simulated phenotype based on the five pre-determined percentage explainability values: {10, 30, 50, 70, 100}, (loosely termed, *heritability* here), which specifies the proportion of phenotypic variance explained by total protein effects. For simplicity, we set the intercept term (or “base” effect) to 0 in all models.

We built random forest using the R *ranger* package and applied the same parameters we use for our real-world dataset analysis (mtry = 500, number of trees = 1500, minimum node size = 5, maximum depth = 82, sqrt(total features)). We analysed the occurrences of parent-child pairs (representing direct conditional relationships between variables), and grandparent-parent-child triplets (representing higher-order interactions). We counted these occurrences across trees to create ranked frequency tables, using dense ranking for ties.

### Interaction Models Considered

The interaction models considered in our synthetic data experiment were based on Wright *et al* ^26^. Complete details of each interaction model is included in Supplementary Table T1. The models considered here are not exhaustive, and some of them have been used primarily to test epistasis in genetics, which may not be entirely realistic for protein interactions.

### Pathway and cluster analysis

Pathway analysis to find enriched pathways was conducted using the R package g:profiler. Network analysis was performed using Cytoscape, using the clusterMaker and STRING Apps. Network analyses was conducted with the “Analyse Network” function in Cytoscape v.2.12. Hub proteins were identified as the nodes with the largest number of edges, as well as highest betweenness centrality. Markov cluster analysis^62^ was performed to find internal clusters between drug classes. The y:files organic edge router visualisation method was used, and nodes were bundled with cytoscape default parameters, except number of handles = 3. Pathway analysis was conducted using the EnrichmentMap plugin.

### Structural modelling

To model our putative binding interactions, we utilised a local implementation of AlphaFold3^63^. Amino acid sequences for EGFR (https://www.uniprot.org/uniprotkb/P00533/entry), ERBB2 (https://www.uniprot.org/uniprotkb/P04626/entry) and ATL3 (https://www.uniprot.org/uniprotkb/Q6DD88/entry) were extracted from UniProt. SMILES structure for lapatinib was extracted from the EMBL-EBI CHEMBL database.

Following structure prediction, potential ligand-binding residues were identified by integrating predicted aligned error (PAE) matrices, directly derived from AlphaFold3 predictions, with residue-level solvent accessibility. A standalone python script was developed to combine AlphaFold3 PAE scores with solvent-accessible surface area (SASA) values calculated using the Shrake–Rupley algorithm (via Bio.PDB.SASA). Each residue was assigned a composite binding likelihood score based on both PAE and SASA, with high-scoring residues identified as candidate ligand-binding sites.

To further evaluate the binding sites predicted by AlphaFold3, we performed a pocket search to complement and strengthen the structural predictions with additional conservation and druggability assessments. The Fpocket^64^ algorithm was applied both to the primary AlphaFold3-predicted structure and to a set of model replicates generated using different random seeds. A custom Python script was then used to analyse the resulting pockets by comparing them across models, quantifying residue-level conservation, and scoring each pocket based on its overlap with highly and moderately conserved ligand-binding residues. Finally, the pockets were ranked using conservation-weighted scores and Fpocket-derived druggability indices.

We estimated the approximate binding affinity of Lapatinib with the proteins using the AutoDock Vina scoring method via the Python API. To begin, we prepared the AlphaFold3-predicted protein–ligand complexes for the calculation. Specifically, we protonated both the protein and ligand at physiological pH (7.4) to ensure chemically accurate hydrogen placement and protonation states, without altering the heavy-atom coordinates, thereby preserving the original geometry predicted by AlphaFold3. Next, we generated receptor maps for Vina scoring, using a docking grid centered on the ligand coordinates and defining a cuboid with dimensions of 20 Å along each principal axis (x, y, z) to encompass the predicted binding site. We first evaluated the raw binding affinity score of the predicted pose.

Subsequently, we performed local energy minimization using Vina’s *minimize()* function to relieve minor steric clashes introduced during hydrogen addition, obtaining a refined binding score. We report both scores for the complexes.

## Compute resources

To run our analyses, we launched a SLURM array job with the following parameters: --ntasks-per-node 1 --cpus-per-task=1 --mem 8gb -t 24:00:00 --array 1-40. Regression trees were built using the R package *ranger* and subsequent analyses were conducted using the python packages pandas, numpy, statsmodels and itertools.

## Tool availability

SynerOmics is available on GitHub https://github.com/Yatish0833/multiomics-synergies.

## Additional files

1. File name: ‘Supplementary_Data_D1.csv’ Title of data: ProCan DepMap Sanger all hits Description of data: Full list of significant (adjusted p-value < 0.01) pairwise synergies for the ProCan-DepMapSanger dataset, for all drugs.
2. File name: ‘Supplementary_Data_D2.csv’ Title of data: Validation datasets all hits Description of data: Full list of significant (adjusted p-value < 0.01) pairwise synergies for the ProCan-DepMapSanger dataset, for all drugs.
3. File name: ‘Supplementary_Data_D3.pdf’ Title of data: Fpocket report EGFR lapatinib Description of data: *Report containing Fpocket-derived druggability and pocket assessment for EGFR*. Ligand-binding pockets were identified and characterized based on geometric and physicochemical descriptors. Druggability scores, pocket volumes, hydrophobicity indices, and polarity metrics were computed to evaluate the potential of each cavity for small-molecule targeting.
4. File name: ‘Supplementary_Data_D4.pdf’ Title of data: Fpocket report ERBB2lapatinib Description of data: *Report containing Fpocket-derived druggability and pocket assessment for ERBB2*. Ligand-binding pockets were identified and characterized based on geometric and physicochemical descriptors. Druggability scores, pocket volumes, hydrophobicity indices, and polarity metrics were computed to evaluate the potential of each cavity for small-molecule targeting.
5. File name: ‘Supplementary_Data Title of data: Fpocket report ATL3 lapatinib Description of data: *Report containing Fpocket-derived druggability and pocket assessment for ATL3*. Ligand-binding pockets were identified and characterized based on geometric and physicochemical descriptors. Druggability scores, pocket volumes, hydrophobicity indices, and polarity metrics were computed to evaluate the potential of each cavity for small-molecule targeting.
6. File name: ‘Supplementary_Figures.docx’ Title of data: Supplementary Figures and Materials S1 to S9 Description of data: Supplementary Figures and Materials S1 to S9
7. File name ‘Supplementary_Tables.xlsx Title of data: Supplementary Tables S1 to S15 Description of data: Supplementary Tables S1 to S15

## Declarations

Ethics approval and consent to participate

Not applicable.

## Consent for publication

Not applicable.

## Availability of data and materials

The datasets supporting the conclusions of this article are available in the Github repository, https://github.com/Yatish0833/multiomics-synergies.

## Authors’ contributions

PRM: conceptualization, methodology, formal analysis, investigation, software, visualization, writing (original draft); AK1: formal analysis, data interpretation; RR, BH: software development; JW, LE, HAM, AK2: data analysis; MK: tool development, data interpretation; LW: supervision; YJ: supervision, software oversight; RR, QZ: project oversight; NT, DB: supervision, writing – review & editing. All authors reviewed and approved the final manuscript. AK1: Anubhav Kaphle, AK2: Anne Klein

## Supplementary Figures and Materials

### S1: Synthetic data experiments

The main goals of the synthetic data experiments are twofold. First, we aim to verify whether variables (proteins, in our case) that are truly (statistically) interacting co-occur more frequently as parent-child nodes along the branches of the regression trees than when they are not. Second, we seek to determine whether the co-occurrence count threshold of greater than 2, used in Syneromics’ first step, provides fair detection of true interactions. This threshold serves as a criterion for screening potentially interacting variables.

#### We conducted the simulation experiments in two steps

In the first step, we simulate our phenotype values (synthetic drug inhibitory concentration, IC_50_) influenced by a single chosen interaction model (Supplementary Figure S0). Initially, we consider both order-2 and order-3 interactions. We call this the simple phenotype simulation setup. The main paper describes this experiment and explains the results. As previously discussed, we were unable to detect any true triplets for the order-3 interaction models in the simple phenotype case. Therefore, we decided to focus solely on order-2 models in our subsequent analysis.

**Supplementary Figure S0:**
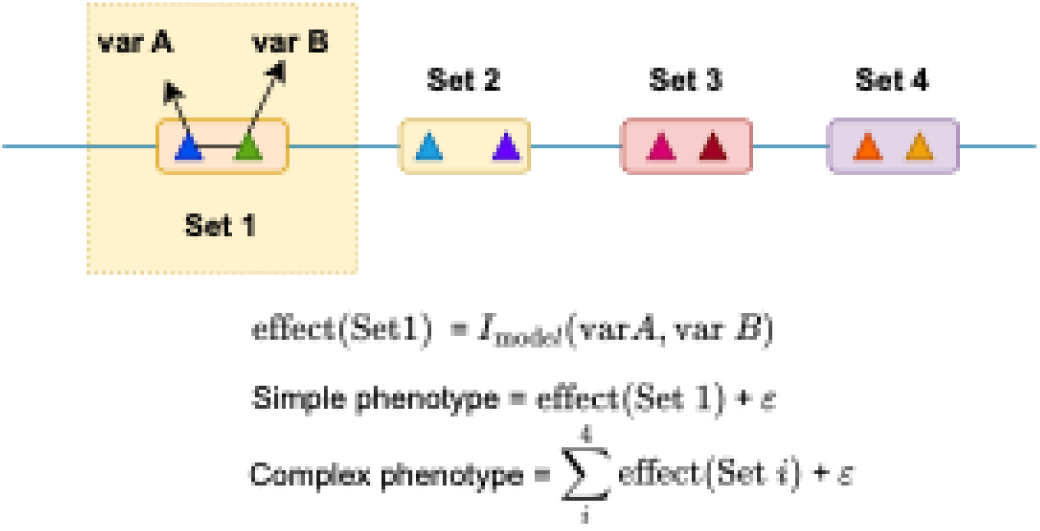
A schematic showing the steps for generating simple and complex phenotypes based on mutually exclusive variable sets, each containing exactly two variables (e.g, var A and var B). Simple phenotypes are influenced by exactly one variable set through a selected interaction model Imodel, whereas complex phenotypes can be influenced by up to four variable sets (shown here is the variable set = 4 case). We add noise ∈ to the generation process by sampling it randomly from a standard Gaussian distribution with its scale adjusted according to the selected heritability value.

In the second step, we consider a complex phenotype setup, where the phenotype is influenced by multiple variable sets (each containing exactly two variables), acting through different interaction models (Supplementary Figure S0). There can be at most four variable sets (including one for completeness), and each set is mutually exclusive, meaning no variable appears in more than one set. Additionally, we ensured that the joint effect of the sets on the complex phenotype follows an additive model.

##### a. Simple phenotype simulation

We discuss our simulation setup for simple phenotype case in detail in the main paper. Briefly, we simulated 10 data versions under each of the 9 considered interaction models, across 5 heritability values. Wherever applicable, we randomly assign marginal effect sizes of variables as either −1 or 1. Interaction effect sizes were randomly sampled from a standard Gaussian distribution with a standard deviation of 1/3, ensuring that interaction effect sizes remain smaller than marginal effect sizes.

To assess co-occurrence frequency, we use the median rank (across 10 data points but excluding 0 when they occur – hence using conditional analysis approach) of the true variable pair among all pairs captured in the decision trees, discussed in Figure 2 of the main paper.

Below, we also provide the median count of occurrences of true variable pairs for each interaction model. We exclude count of 0 for median calculation as that corresponds to no detection due to the stochasticity involved in tree-building process. Thus, the results represent a conditional performance evaluation, where aggregate measure only convey information when true pairs are detected. We take into consideration count of 0 for computing overall detection rate, which is discussed below.

**Supplementary Figure S1:**
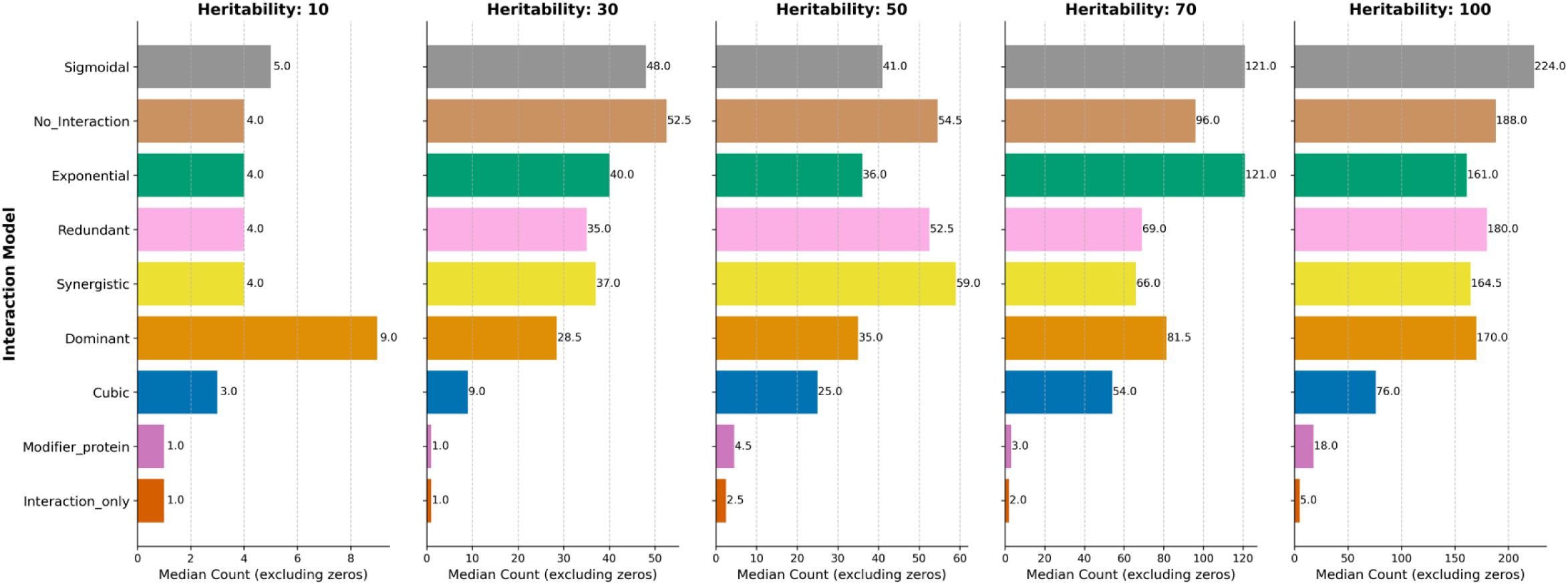
Median co-occurrence counts for true variable pairs across heritability values for different interaction models considered in the synthetic data experiments. The median count excludes 0 (no detection) from the calculation.

To assess whether count >= 2 is a reasonable threshold for identifying true variables, we also calculate the rate of true detection rate for that count threshold. The rate of detection is simply the percentage of successful identifications of true variable pairs for a given interaction model at a chosen heritability value when using count >= 2 for screening. We also calculate the heritability-weighted overall detection rate, which can be interpreted as weighted average of detection average with more emphasis on the challenge of low heritability detection. We have plotted the detection rate trajectory below and noted the overall detection rate.

**Supplementary Figure S2:**
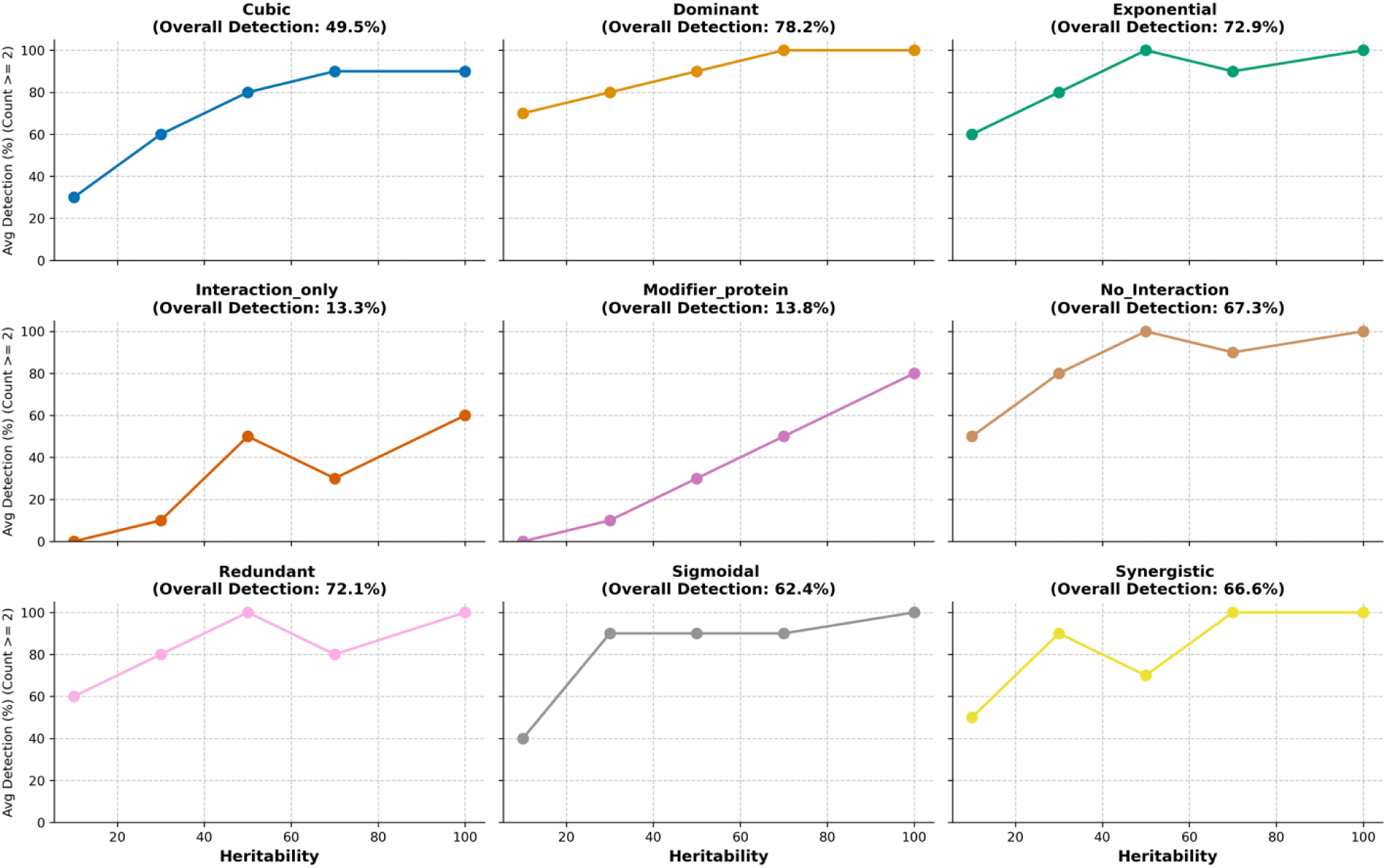
True variable pairs detection rate (%) trajectory for each interaction model across the heritability spectrum. The overall detection is calculated based on a weighted average of the detection rates across heritability values, emphasising larger weights for successful detection at lower heritability values.

There is an upward trend in the detection rate with increasing heritability values, as expected (Supplementary Figure S2). With higher heritability, it becomes easier to detect the interaction signal. When weighted based on heritability, the average overall detection rate is highest for pairs acting through the dominant model (78.2%) and lowest for the interaction-only model (13.3%). The synergistic model, which we particularly focus on in our actual proteomic analysis, has a decent overall detection rate of 66.6% across the heritability spectrum, however, has higher detection rates at higher heritability values (0.7, 1.0) than other models.

##### b. Complex phenotype simulation

We simulate complex phenotypes using the same parameter setup as in the simple phenotype case. Briefly, we randomly assign marginal effect size to either −1 or 1, and interaction effect size are drawn from a standard Gaussian with standard 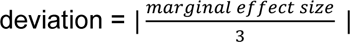

For each variable set, we calculate its effect on the phenotype based on an interaction model randomly selected from the 9 considered models.

In each iteration of complex phenotype generation, we randomly determine the number of variable sets between 1 and 4. Each set is then assigned a randomly chosen interaction model. The two variables in each set are also randomly selected, ensuring that their Pearson’ s correlation (here based on protein expression values) with any other variable in the dataset does not exceed 0.7. We then sum the effects from the variable sets. Finally, we add noise to the final effects by sampling from a standard Gaussian distribution, with standard deviation determined by the heritability values used in the simulation, and to generate ‘noisy’ complex phenotype. We simulate a total of 100 data replicates for each heritability value.

Since variable sets and interaction models are selected randomly, we don’t expect each to have same number of data points for each interaction model across heritability and variable set combination.

**Supplementary Figure S3:**
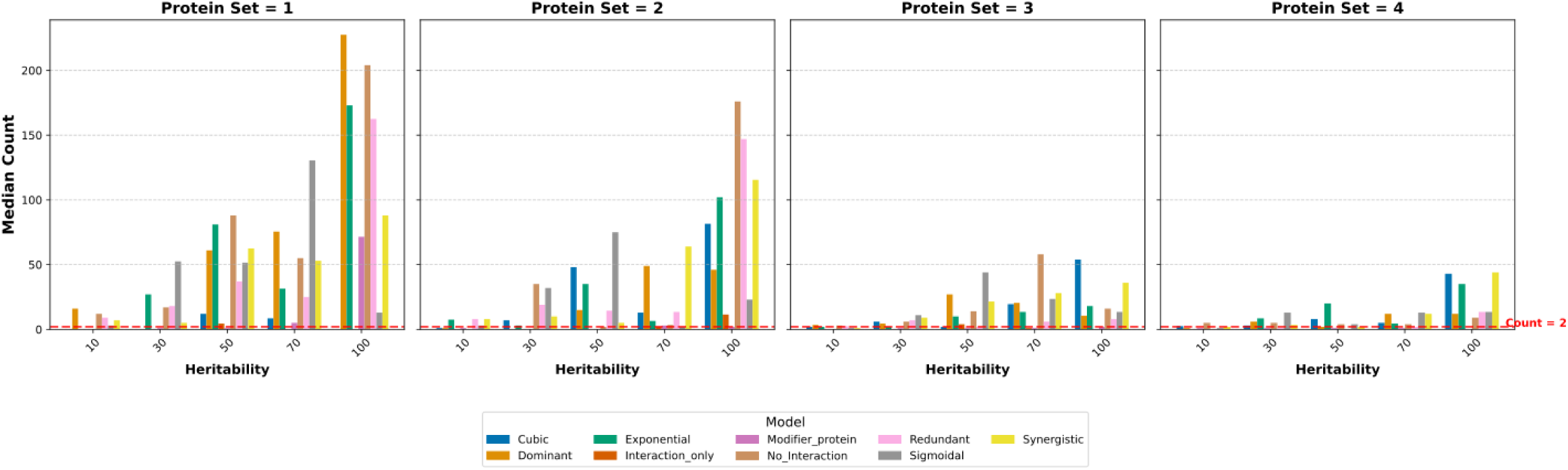
Median (excluding zeros) co-occurrences counts for true variable pairs across variable sets, and heritability values for the 9 interaction models considered in the simulation.

As expected, we observe an upward trend in the median count with increasing heritability values across variable sets (Supplementary Figure S3). We also observe an overall attenuation in the median count as the number of variable sets increases at any heritability value, highlighting the effect of increasing complexity. Notably, even as complexity increases, some variables - acting through interaction models such as cubic, synergistic, exponential, sigmoidal – are still detected at modest heritability levels, meeting the count>=2 threshold. We formally assess the detection rate below.

To obtain the mean detection rate for a given interaction model, we first compute the weighted harmonic mean of the detection rate for a given heritability value across variable sets. We use the number of variable sets as our weight to place greater emphasis on detections when multiple variable sets influence phenotype, which represent the more challenging cases. We then compute the weighted average of these harmonic means to obtain the overall (aggregate) detection rate, using the inverse of heritability values as weights to emphasize detection in lower heritability ranges during mean calculation. The overall detection rate reflects the effectiveness of using the threshold of >=2 to screen for potentially interacting pairs using random forest models, across different complexities and heritability levels, with a focus on harder-to-detect pairs in a low heritability and high complexity setting.

As expected, we see an upward trajectory in the detection rate, although with weaker slopes compared to the simple phenotype setup, reflecting the challenges of increased complexity. The overall detection rate has also fallen by about 50% across interaction models compared to the previous setup. The dominant interaction model is no longer the best detected in this complex setting. We observe fairly improved detection for synergistic models compared to other models, even under challenging conditions (high complexity and low heritability). Interaction_only and Modifier_protein have low overall detection rates (3.40% and 9.65%), suggesting that the random forest (with a count threshold of ≥2) struggles to capture true interactions in the challenging conditions.

**Supplementary Figure S4:**
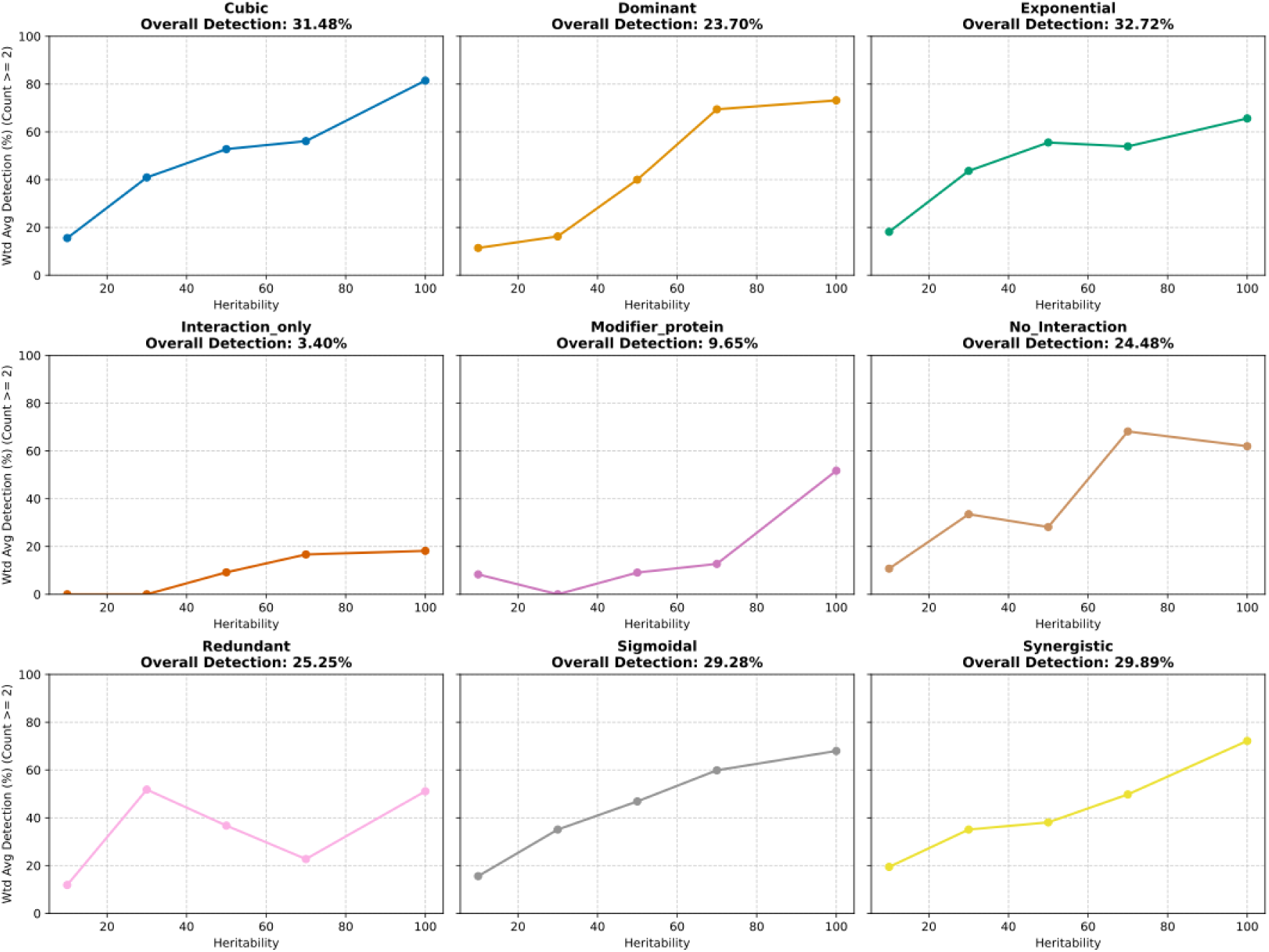
The trajectory of the true variable pairs detection rate (%) for each interaction model across the variable sets and heritability spectrum. We calculate the overall detection rate by first taking the weighted harmonic mean of the detection rate across protein sets, giving more importance to detection in larger protein sets. We then compute the weighted average of these harmonic means across heritability values to obtain the overall detection rate.

**Supplementary Figure S5:**
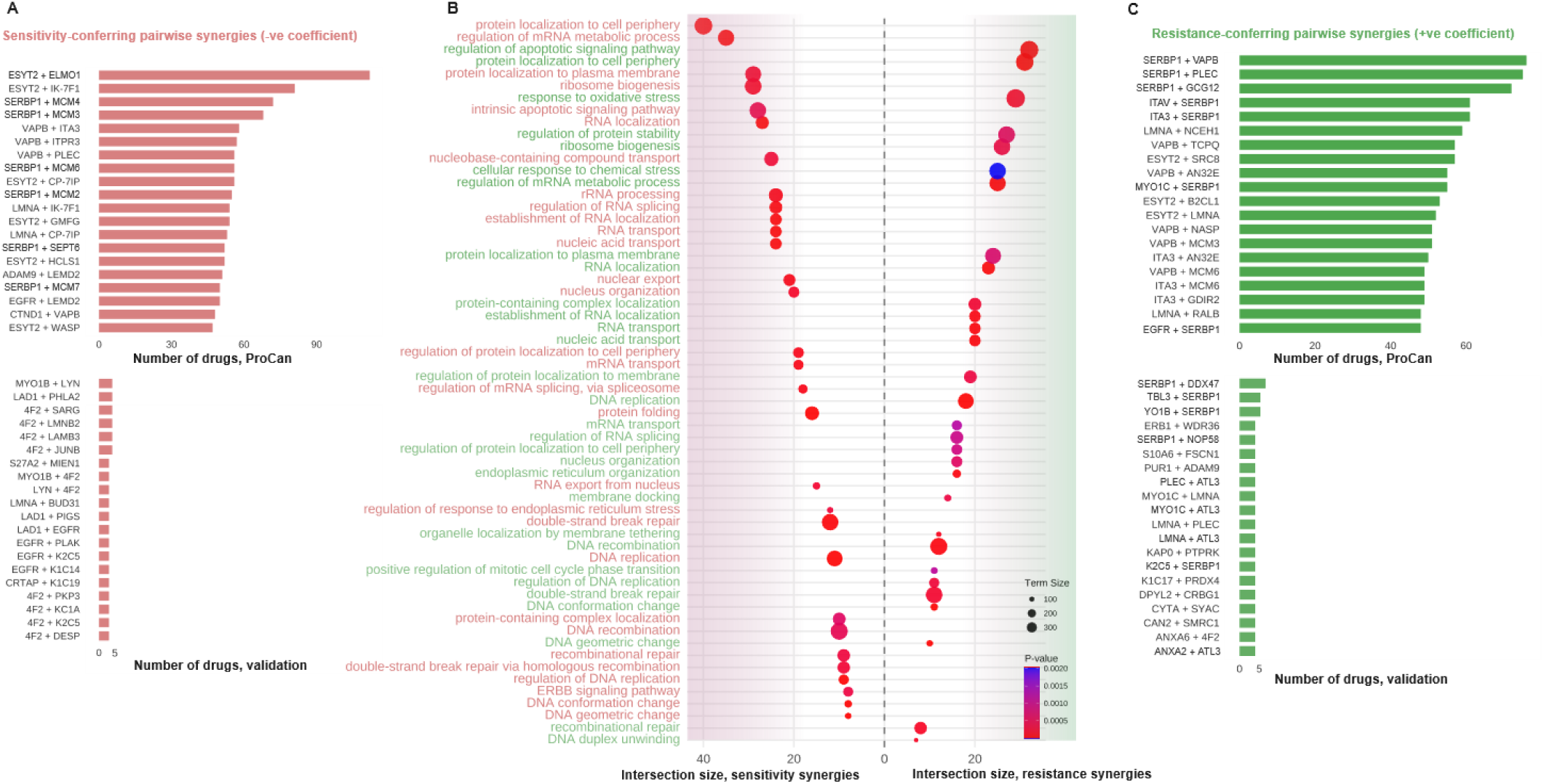
Pathway analysis of shared synergies. goProfiler analysis of shared synergies in both datasets, coloured by enrichment for resistance synergies (green text) and sensitivity synergies (red text). Pathways to the left of the dotted line represent pathway enrichment of sensitivity synergies, while pathways to the left of the dotted line represent pathway enrichment for resistance synergies. Circle sizes represent overall size of the pathway term, and colour represents p-value.

**Supplementary Figure S6:**
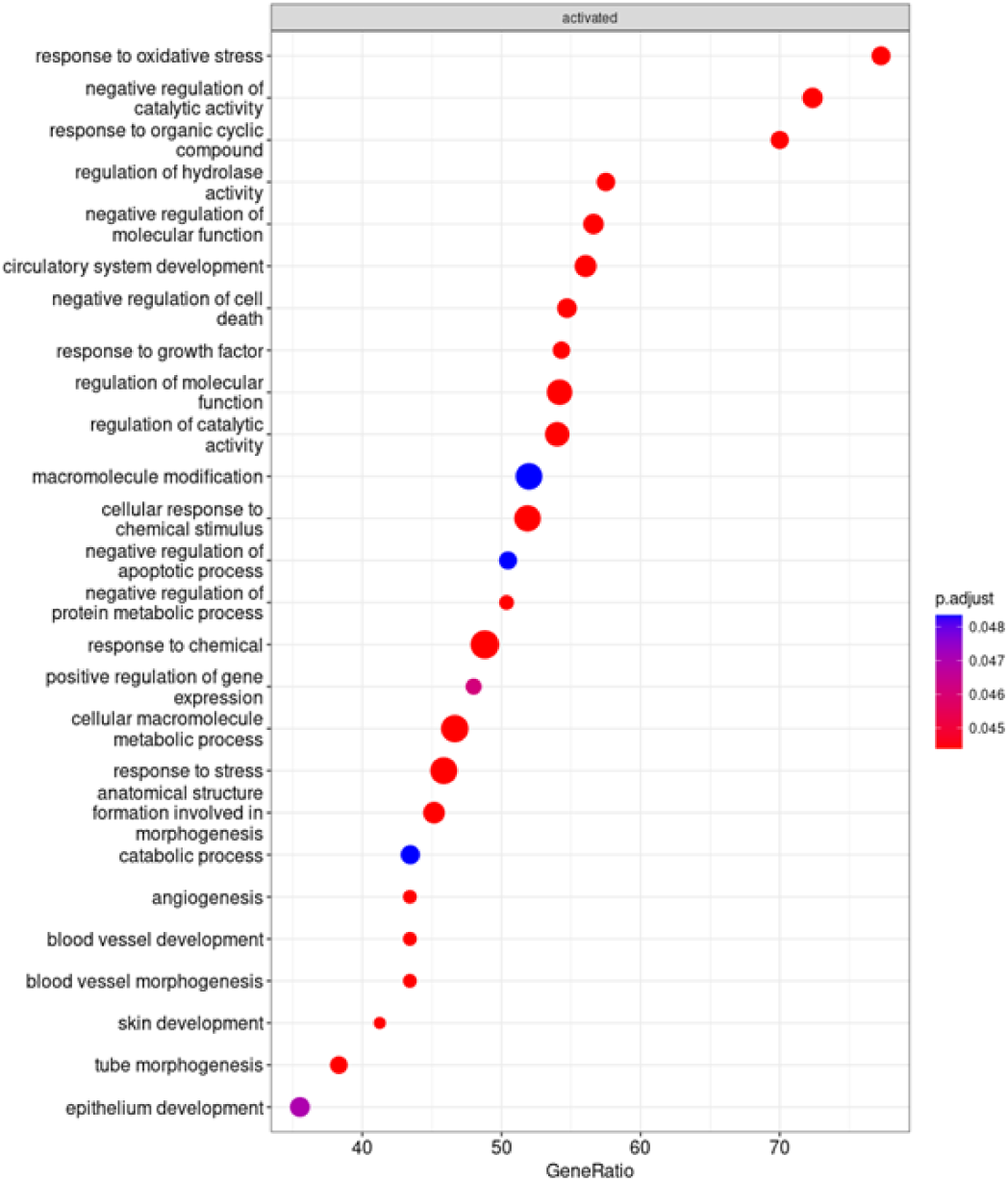
GoProfiler pathway analysis of significant interactions associated with drug susceptibility. GeneRatio represents the number of proteins in the set that overlap with our proteins of interest.

**Supplementary Figure S7:**
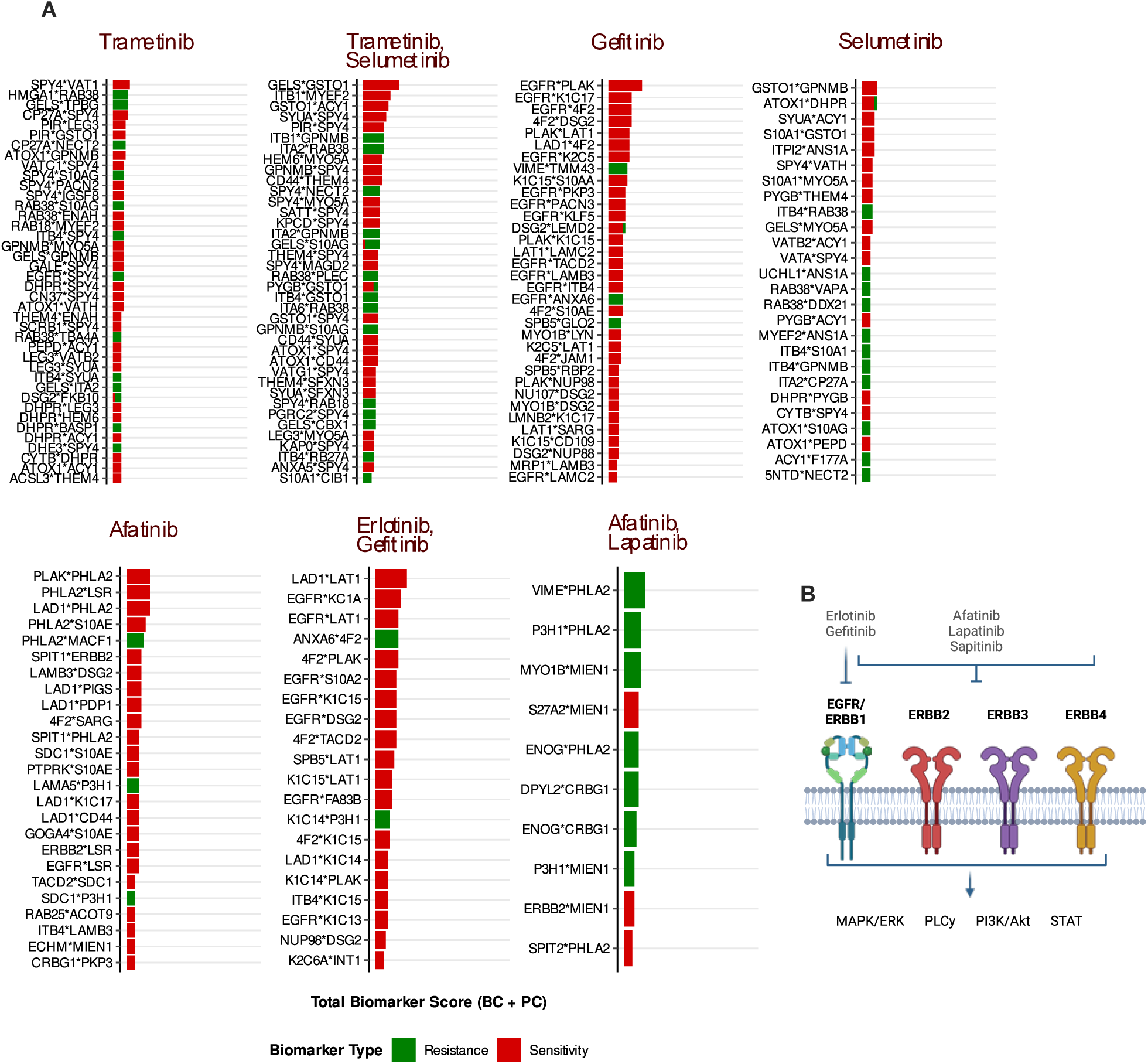
Top resistance and sensitivity pairwise synergies replicated across both ProCan and validation datasets. B, ERBB family receptors and targeting drugs (adapted from Schramm et al.55).

**Supplementary Figure S8:**
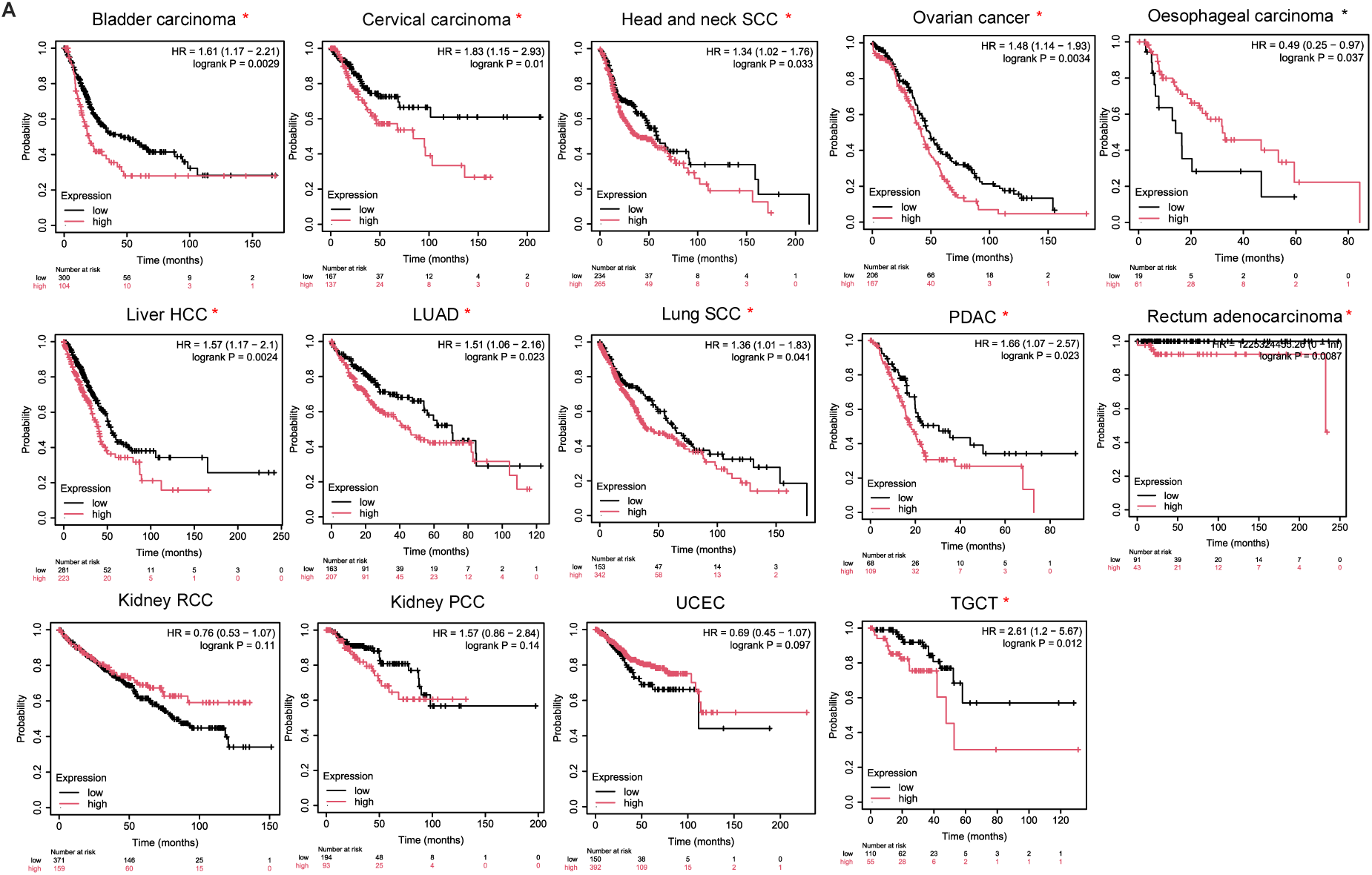
KM Plotter analysis survival analysis of TCGA mRNA datasets using gene signature (mean expression) of ATL3, ESYT2, MYO1C, LMNA and PLEC. Red asterisk * represents a significant association between worse survival and high expression of gene signature, black asterisk * represents a significant association between better survival and high expression of gene signature.

**Supplementary Figure S9:**
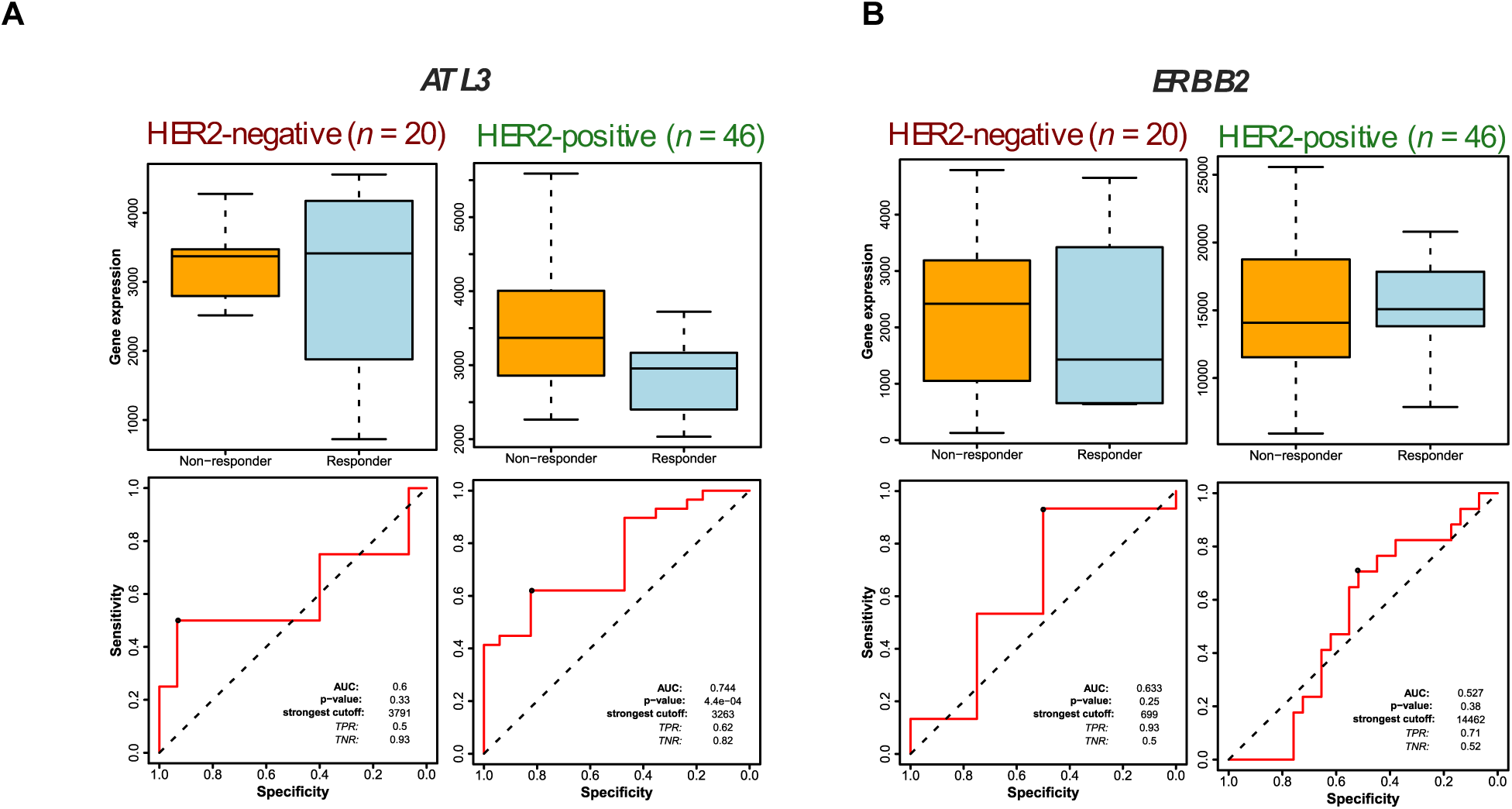
ROCplot analysis comparing predictive power of ATL3 vs. ERBB2 for lapatinib resistance

